# RNA interference dynamics in juvenile *Fasciola hepatica* are altered during *in vitro* growth and development

**DOI:** 10.1101/2020.06.06.137646

**Authors:** Paul McCusker, Wasim Hussain, Paul McVeigh, Erin McCammick, Nathan G. Clarke, Emily Robb, Fiona M. McKay, Peter M. Brophy, David J. Timson, Angela Mousley, Nikki J. Marks, Aaron G. Maule

**Affiliations:** Microbe and Pathogen Biology, Institute for Global Food Security, School of Biological Sciences, Queen’s University Belfast, Belfast, UK; Institute of Biological, Environmental & Rural Sciences, Aberystwyth University, Aberystwyth, UK; School of Pharmacy & Biomolecular Sciences, University of Brighton, Brighton, UK

**Keywords:** *Fasciola*, Liver fluke, RNAi, Argonaute, Calmodulin, σGST

## Abstract

For over a decade RNA interference (RNAi) has been an important molecular tool for functional genomics studies in parasitic flatworms. Despite this, our understanding of RNAi dynamics in many flatworm parasites, such as the temperate liver fluke (*Fasciola hepatica*), remains rudimentary. The ability to maintain developing juvenile fluke *in vitro* provides the opportunity to perform functional studies during development of the key pathogenic life stage. Here, we investigate the RNAi competence of developing juvenile liver fluke. Firstly, all life stages examined possess, and express, core candidate RNAi effectors encouraging the hypothesis that all life stages of *F. hepatica* are RNAi competent. RNAi effector analyses supported growing evidence that parasitic flatworms have evolved a separate clade of RNAi effectors with unknown function. Secondly, we assessed the impact of growth / development during *in vitro* culture on RNAi in *F. hepatica* juveniles and found that during the first week post-excystment liver fluke juveniles exhibit quantitatively lower RNAi mediated transcript knockdown when maintained in growth inducing media. This did not appear to occur in older *in vitro* juveniles, suggesting that rapidly shifting transcript dynamics over the first week following excystment alters RNAi efficacy after a single 24 hour exposure to double stranded (ds)RNA. Finally, RNAi efficiency was found to be improved through use of a repeated dsRNA exposure methodology that has facilitated silencing of genes in a range of tissues, thereby increasing the utility of RNAi as a functional genomics tool in *F. hepatica*.

## 1. Introduction

Fasciolosis, caused by liver fluke of the genus *Fasciola*, is responsible for considerable global agri-food sector losses (de Waal et al., 2007; Gray et al., 2008; Adrien et al., 2013; Khoramian et al., 2014) and, is recognised as a neglected tropical disease by the World Health Organisation (WHO, 2015). Treatment for *Fasciola* infection relies on a small cohort of anthelmintics, although triclabendazole remains the current drug of choice due to its efficacy against most life stages in the mammalian host (Kelley et al., 2016). However, field resistance to triclabendazole was first reported in the mid-1990s (Overend and Bowen, 1995), and since then resistance cases have been reported worldwide (Kelley et al., 2016). Due to increasing resistance to triclabendazole and other flukicides (Novobilský and Höglund, 2015) it is critical that either (i) the current drug cohort is improved, or (ii) novel drug/vaccine targets are identified and utilized to drive the discovery/development of new flukicides or a fluke vaccine.

RNA interference (RNAi) is an amenable functional genomics tool that could prove pivotal in the discovery and validation of new flukicide and vaccine targets. Since its discovery (Fire et al., 1998), RNAi has facilitated reverse genetic screens in many parasitic helminths. Although there has been limited success with RNAi in animal parasitic nematodes (Viney and Thompson, 2008), RNAi has been successfully performed in a range of parasitic platyhelminths, such as *Schistosoma, Fasciola, Moniezia, Echinococcus, Opisthorchis, Hymenolepis* and *Clonorchis* (Boyle et al., 2003; Skelly et al., 2003; McGonigle et al., 2008; Pierson et al., 2010; Spiliotis et al., 2010; Sripa et al., 2011; Pouchkina-Stantcheva et al., 2013; Wang et al., 2014). RNAi in schistosomes has resulted in a variety of quantifiable phenotypes, informing the function and roles of many genes in this important pathogen. RNAi-based therapies have also been touted, although only one study demonstrated a modest reduction in infection following the injection of schistosome-targeting dsRNA into schistosome-infected mice (Pereira et al., 2008).

Five studies on *in vitro* RNAi in *F. hepatica* juveniles have been published (McGonigle et al., 2008; Rinaldi et al., 2008; Dell’Oca et al., 2014; McVeigh et al., 2014; McCammick et al., 2016). The cathepsin silencing reported by McGonigle et al. (2008) described reduced motility and aberrant migration through the intestinal wall whilst McCammick et al. (2016) reported a reduction in juvenile fluke growth and hyperactivity following the knockdown of calmodulin genes. Silencing of the exogenous gene firefly luciferase resulted in a reduction in luciferase activity in newly excysted juveniles (NEJs) (Rinaldi et al. 2008) and various other genes have been silenced without any phenotypes being reported (Dell’Oca et al. 2014; McVeigh et al., 2014). To date, there have been no reports of serious morphological damage or lethality following RNAi studies in juvenile *F. hepatica* and it is unclear if this is simply due to the gene complement silenced thus far, is due to the *in vitro* culture conditions or to RNAi pathway dynamics in early stage *F. hepatica*.

In an effort to better understand the biology underpinning RNAi in liver fluke, we exploited available genomic / transcriptomic datasets for fluke to build a better understanding of the panel of RNAi effectors employed by these worms. Previous studies on the RNAi dynamics of schistosomes and *Fasciola* have focused on optimizing the delivery of dsRNA / siRNA to improve RNAi penetrance and persistence (Ndegwa et al., 2007; Stefanić et al., 2010; Dell’Oca et al., 2014; McVeigh et al., 2014), but this study focuses on the impacts of growth / development on RNAi effectiveness. The latter is possible because we recently developed an *in vitro* maintenance and growth platform (McCusker et al., 2016) that allows the sustained maintenance of growing juvenile fluke. This platform enables the examination of RNAi responses in juvenile worms across early-stages of development in the mammalian host and over sustained timeframes, allowing us to explore the biology of growing fluke and to optimize RNAi-based functional genomics.

## 2. Methods

### 2.1 RNAi effector identification

Protein sequences of RNAi / miRNA pathway effectors from 21 species, both invertebrates and vertebrates, were recovered from www.wormbase.org, parasite.wormbase.org, www.ncbi.nlm.nih.gov/protein, www.genedb.org and Dalzell et al. (2011) (Supplementary File 1). Family groups of RNAi effectors were aligned using Clustal Omega (www.ebi.ac.uk/Tools/msa/clustalo/) and converted to Stockholm format. Gblocks (molevol.cmima.csic.es/castresana/Gblocks_server.html) was used to identify conserved regions, which were then used to assemble query sequences on HMMER software (www.hmmer.org/). Query sequences were used to examine a *F. hepatica* genome predicted protein dataset (Cwiklinski et al., 2015), via a Hidden Markov Model (HMM) approach. There were 103 individual hits identified that were used in BLASTp searches against non-redundant databases to confirm sequence identity with conserved regions identified through Interpro (www.ebi.ac.uk/interpro/search/sequence-search). Note that initial RNAi effector protein identification and primer design was performed on sequences from an older genome assembly. An updated assembly was released whilst this manuscript was being prepared. We have used our original results to identify the most recent accession numbers (found on http://parasite.wormbase.org) using BLASTp searches.

Protein model prediction was performed using Protein Homology / analogY Recognition Engine V 2.0 (Kelley et al., 2015). Putative effectors were aligned with *Homo sapiens, Drosophila melanogaster, Caenorhabditis elegans, Schistosoma mansoni* and *Schistosoma japonicum* orthologues using MUSCLE (www.ebi.ac.uk/Tools/msa/muscle/), and visualised in Jalview (Waterhouse et al., 2009). Maximum-likelihood trees were constructed in MEGA7 (www.megasoftware.net/) by performing 1000 bootstrap replications.

### 2.2 Excystment and culture

UK isolate *F. hepatica* eggs and metacercariae (from University of Liverpool, UK and Queen’s University Belfast, UK) were used for expression analyses, whilst *F. hepatica* Italian isolate metacercariae (Ridgeway Research, UK) and North-Western USA metacercariae (Baldwin Aquatics, USA) were used in RNAi trials (these isolates did not differ in RNAi susceptibility). Outer cyst walls were removed through mechanical separation of outer walls with a scalpel prior to incubation in 10% thin bleach at RT (5 min). Metacercariae were rinsed in dH_2_O five times before excystments were carried out as described in McVeigh et al. (2014). Newly excysted juveniles (NEJs; excysted within 4 h), were transferred to RPMI (RPMI 1640 (ThermoFisher Scientific) with Antibiotic/Antimycotic Solution x100 (Sigma-Aldrich) at 1:100). Worms for culture (referred to as juveniles) were placed in 96-well round bottom plates containing either RPMI or CS50 (50% chicken serum (Sigma-Aldrich) in RPMI).

### 2.3 Adult F. hepatica recovery

Adult *F. hepatica* were recovered from condemned sheep livers at a local abattoir (ABP Meats, Lurgan, Northern Ireland), and transported to the laboratory in 15 mM NaCl (37°C). Fluke were maintained at 37°C in 15 mM NaCl (replaced every 30 min) for around 1.5 h to encourage the regurgitation of gut contents.

### 2.4 RNAi effector expression

Eggs (1000 / replicate), cysts (100 / replicate), NEJs (100 / replicate), two-week-old juveniles (20 / replicate) and adults (1 / replicate) were snap frozen in liquid nitrogen before lysis and extraction using the Dynabeads mRNA DIRECT Kit (ThermoFisher Scientific). Eluted mRNA samples were treated with TURBO DNase (ThermoFisher Scientific) prior to cDNA synthesis (High Capacity mRNA to cDNA Kit (ThermoFisher Scientific)).

Primers (Metabion International AG; Supplementary Table 1) were designed using Primer3Plus (Untergasser et al., 2007). End-point PCRs were used to confirm expression of RNAi effectors identified through HMM profiling in eggs, cysts, NEJs, two-week-old juveniles grown in CS50 and adults using the FastStart™ Taq DNA Polymerase, dNTPack (Sigma-Aldrich) with 0.4 µM of each primer, and 1 µl 1:1 diluted cDNA. PCR products were size confirmed on a 0.1% agarose gel and then sequence confirmed (Eurofins Scientific) after PCR clean up (ChargeSwitch PCR Clean-Up Kit (ThermoFisher Scientific)).

### 2.5 RNAi effector qPCR

RNAi effector expression was compared between worms maintained in RPMI and CS50 by extracting mRNA from 20 worms / replicate after one week as described above. RNA concentrations were normalised using the Qubit^®^ RNA High Sensitivity Assay Kit (ThermoFisher Scientific) prior to reverse transcription (as above). qPCRs were run using 2 µl cDNA (diluted 1:1) in a 10 µl reaction with 5 µl SensiFAST SBYR No-ROX Kit (Bioline) and 0.1 µM of each primer. ΔCt values were obtained using half of the Pfaffl equation (Pfaffl, 2001) with values transformed, via log_2_, to normalise expression.

### 2.6 Juvenile RNAi

Double stranded RNAs (dsRNA) specific to bacterial neomycin (bacterial neomycin phosphotransferase, U55762), *F. hepatica* σ-glutathione-S-transferase (*fhσgst*, DQ974116), and *F. hepatica* calmodulin 2 (*fhcam2*, AM412547) were generated using the T7 RiboMAX™ Express RNAi System (Promega) with primers listed in Supplementary Table 2. Juveniles were excysted, as described above, and transferred into 96-well round bottom plates (20 juveniles / replicate). RNAi trials were run over a period of three days to four weeks with various treatment regimens. Juveniles grown in CS50 were washed three times in 200 µl prior to dsRNA treatment. At the conclusion of trials juveniles were snap frozen prior to mRNA extraction and reverse transcription, as described above. Transcript abundance was measured via qPCR as described above (except with 0.2 µM of each primer) with glyceraldehyde 3-phosphate dehydrogenase (*fhgapdh*, AY005475) used as reference gene. Primers listed in Supplementary Table 2. ΔΔCt values were calculated using the Pfaffl equation, where untreated groups provided control values and dsRNA treated groups (both control dsRNA and target dsRNAs) provided experimental values, before conversion to percentage expression.

Protein knockdown was assessed via western blot as described in McVeigh et al. (2014). Samples were normalised to 2 µg/µl in radioimmunoprecipitation assay (RIPA) buffer using the Qubit^®^ Protein Assay Kit (ThermoFisher Scientific) prior to processing as described in McVeigh et al. (2014) – primary and secondary antisera were as used in McVeigh et al. (2014) and McCammick et al. (2016). Membranes were developed for ∼30 s before washing thoroughly in distilled water. Membranes were dried and scanned to quantify band densities in ImageJ (http://rsbweb.nih.gov/ij/). Band densities were normalised to intensity of control band (FhCaM2 or FhσGST) to allow for relative quantitation. Untreated samples were considered to exhibit mean normal protein expression (100%) to allow control and target dsRNA samples to be expressed as percentages relative to the untreated worms.

### 2.7 Statistical analyses

T-tests and ANOVAs were used to test for significant differences between treatment groups. Graphs were created, and statistical tests carried out using GraphPad Prism 6 for Windows (GraphPad Software, La Jolla California USA, www.graphpad.com).

## 3. Results

### 3.1 RNAi effector identification and expression

Of the 103 genes identified in the HMM screen, 27 were putative RNAi / miRNA pathway effectors of *F. hepatica*. These 27 genes / proteins comprise all of the core machinery believed to be necessary for a functional RNAi pathway (Figure 1A; Supplementary Table 3). Consistent with other parasitic flatworms we found that *F. hepatica* possessed an Argonaute (AGO) that appears to belong to a parasitic flatworm specific AGO clade (Figure 1C), whilst also possessing an AGO from the ‘classic’ clade 1 group (Skinner et al., 2014). This expansion of the AGO family in parasitic flatworms extends to other proteins in the RNA-induced silencing complex (RISC), such as the Fragile X Retardation Syndrome 1 protein (FMR1), with four found in *F. hepatica*, and three in *S. mansoni*. The number of FMR1s varies between species, although most species do not have the three / four seen in the parasitic flatworms. For example, *D. melanogaster* possesses one, whilst mammals have an additional two paralogues, FXR1 and FXR2 (Wan et al., 2000). The parasitic flatworm FMR1s appear to group into three distinct clades with one clade comprised entirely of trematode-only members (Supplementary Figure 1M). These duplications may have occurred very early in the ancestry of parasitic flatworms as they display little similarity to each other, making it difficult to ascertain which FMR1 is the ancestral homologue.

**Figure 1:**
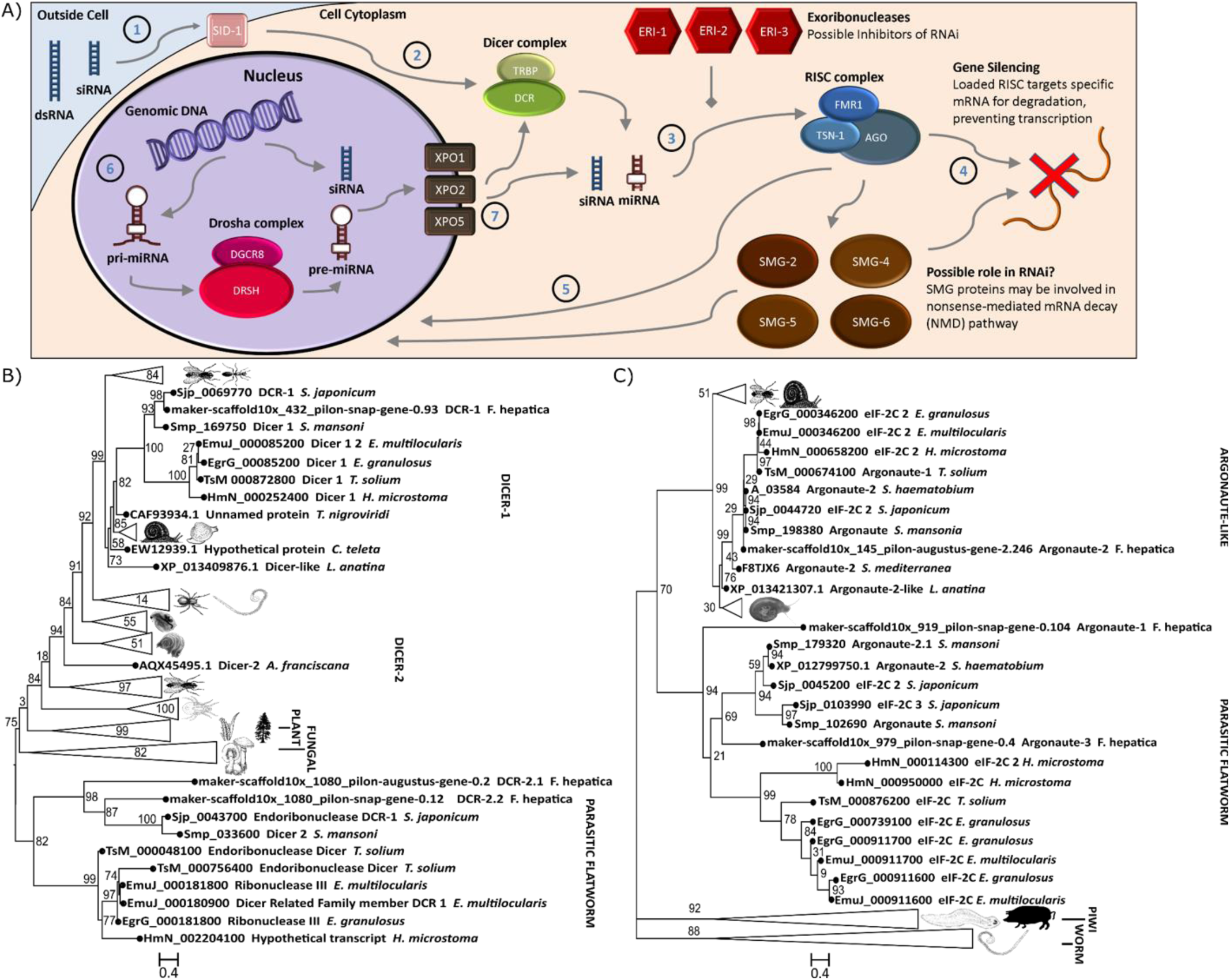
The RNA interference (RNAi) / micro (mi)RNA pathway effectors of *Fasciola hepatica*. A – Summary of the putative RNAi / miRNA pathway in *F. hepatica* where double stranded (ds)RNA enters cells via SID-1 (1). Both dsRNAs and primary (pri)-miRNAs are processed by the dicer complex into short interfering (si)RNA or miRNA (2), which are then incorporated into the RNA-induced silencing complex (RISC) (3) and serve as guide RNAs to target complimentary mRNA for degradation (4). siRNAs and miRNAs may be trafficked into the nucleus (5). In the nucleus the Drosha complex processes pri-miRNAs into pre-miRNAs (6) which are transported out of the nucleus via exportins (7). B – Maximum-likelihood tree of dicer (DCR) proteins in various species reveals distinct evolutionary outgroups in the cestodes and trematodes. C – Maximum-likelihood tree of Argonaute genes in various species reveals loss of (piwi) piRNA processing Argonautes (AGOs), but the presence of a novel AGO clade in parasitic flatworms. Numbers at branch points indicate bootstrap values.

This expansion of RNAi effectors is not limited to RISC forming proteins as parasitic flatworms also have a Dicer-1 (DCR-1) clade that appears to be distinct from other species (Figure 1B). Cestodes were known to have an additional DCR-1 and schistosomes were reported to have an additional DCR-2, a situation now mirrored in *F. hepatica*, suggesting that this duplication is common to trematodes. Although *F. hepatica* DCR-2s are smaller than FhDCR-1 (865 / 1154 vs 2519 amino acids), they still possess the conserved motifs required for RNA binding and nuclease activity (Supplementary Figure 2D).

*F. hepatica*, along with other trematode species appear to possess evolutionarily distinct ERI-1 and ERI-2 genes. *F. hepatica* also possesses two ERI-3 isoforms with both expressed in all life stages examined here (Supplementary Figure 3T and U); note that FhERI-3.2 is considerably smaller (85 vs 211 amino acids) as it is missing a conserved nuclease motif (HD as opposed to DEDHD), although this could be linked to sequence assembly error.

With the exception of *fhtsn-1.2* (maker-scaffold10x_1598_pilon-snap-gene-0.1), the expression of all *F. hepatica* putative RNAi effectors identified through our *in silico* analysis was confirmed in at least one life stage (Supplementary Figure 3). However, RNA-Seq data from Cwiklinski et al. (2015) suggest that *fhtsn-1.2* is also expressed in all life stages (sequence length restricted primer design). *fheri-1* was not detected in embryonated eggs, although it was detected in unembryonated eggs suggesting that RNAi should be possible in even the earliest life stages of *F. hepatica* as well as all others studied here (embryonated eggs, NEJs, two-week-old *in vitro* juveniles and adults).

### 3.2 Juvenile F. hepatica RNAi dynamics

Having concluded that CS50 is currently the most effective maintenance medium for *in vitro* culture (McCusker et al. 2016) we used it to investigate the effect of growth on RNAi responses in juvenile *F. hepatica* to establish the utility of RNAi as a functional genomics tool during *in vitro* development. However, we found that growth in CS50 significantly altered RNAi responses over the first seven days post excystment, with juveniles maintained in RPMI exhibiting greater transcript knockdown than those in CS50 (Figure 2A; *fhcam2* transcript reduction over seven days in RPMI: 86.6%±1.1; in CS50: 66.2%±3.7; *fhσgst* transcript reduction in RPMI: 78.2%±3.6; in CS50: 40.8%±7.3). This transcript knockdown over seven days was less than that after four days of maintenance in CS50 (*fhcam2* transcript reduction in CS50 over four days: 84.8%±2.4; over seven days: 66.2%±3.7 – t-test P=0.0138; *fhσgst* transcript reduction in CS50 over four days: 86.4%±0.6; over seven days: 40.8%±7.3 – t-test P=0.0079)), pointing to transcript recovery for both gene targets, between 4-7 days post dsRNA exposure (Figure 2B).

**Figure 2:**
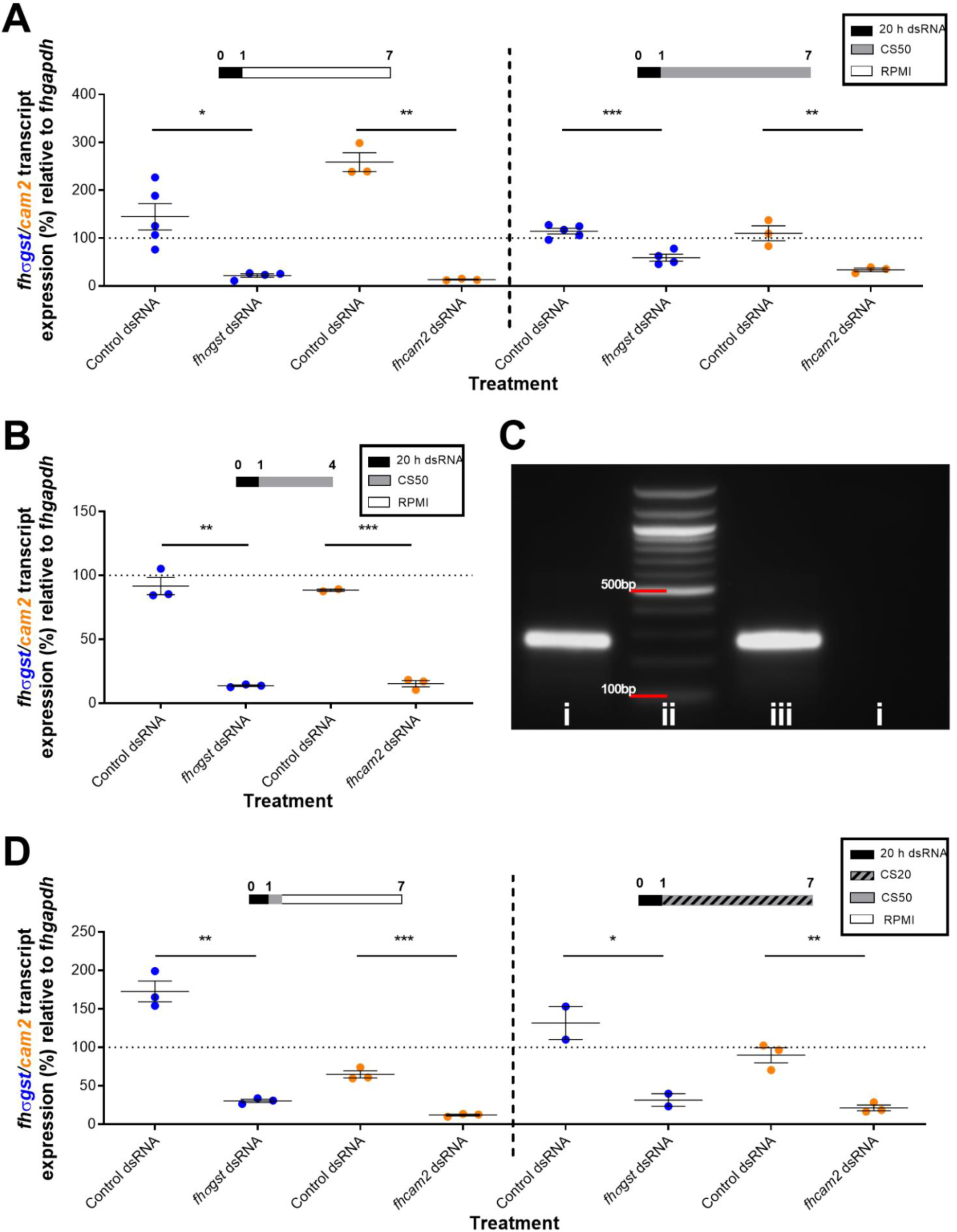
Growth of juvenile *Fasciola hepatica* over the first week following excystment alters RNA interference (RNAi) dynamics. A – Percentage expression of *F. hepatica* sigma glutathione transferase (*fhσgst*) / *F. hepatica* calmodulin 2 (*fhcam2*) transcript relative to *F. hepatica* glyceraldehyde phosphate dehydrogenase (*fhgapdh*) after double stranded RNA (dsRNA) exposure and six days maintenance in 50% chicken serum (CS50) or RPMI shows reduced knockdown in juveniles maintained in CS50. B – Percentage expression of *fhσgst* / *fhcam2* transcript relative to *fhgapdh* after dsRNA exposure and three days maintenance in CS50 shows robust knockdown for both genes. C – Agarose gel electrophoresis image showing (i) 100 ng/µl of dsRNA in RPMI, (ii) 100 bp ladder (iii) 100 ng/µl dsRNA in RPMI at 37°C for 2 h and, (iv) 100 ng/µl dsRNA in CS50 at 37°C for 2 h, showing dsRNA degradation in CS50. D – Percentage expression of *fhσgst* / *fhcam2* transcript relative to *fhgapdh* after dsRNA exposure followed by either dsRNA removal or six days maintenance in 20% chicken serum (CS20) showing how dsRNA removal has little impact on knockdown. Treatment timelines (in days) above graphs show media juveniles were maintained in; y axis dotted line at 100% represents mean transcript expression in untreated juveniles; colours of gene in y axis correspond to gene expression on graphs. Unpaired t-tests used to test for significant differences where *, P<0.05; **, p<0.01; ***, p<0.001; ****, p<0.0001.

Transcript recovery could have been caused by either the degradation of residual dsRNA or growth-dependent transcript dynamics, or a combination of both. To test the importance of residual dsRNA in sustaining knockdown, CS50 (which has endogenous nuclease activity; Kawaguchi et al., 2009) was used to remove / degrade dsRNA (Figure 2C) over two hours following dsRNA soak, before the CS50 was washed out and replaced with RPMI. This induced knockdown comparable to that seen in juveniles maintained continually in RPMI for both genes (Figure 2D; transcript reduction *fhcam2* in RPMI: 86.6%±1.1; in degraded dsRNA: 87.9%±1.1 – t-test: P=0.597; *fhσgst* in RPMI: 78.2%±3.6; in degraded dsRNA: 69.5%±2.1 – t-test: P=0.452), suggesting that reduction in RNAi-mediated knockdown was not linked to residual dsRNA degradation. To test the role of growth we maintained juveniles in CS20 (instead of CS50) following dsRNA exposure as CS20 induces a slower growth rate over one week (McCusker et al. 2016). Knockdown under these conditions lay between that seen in RPMI and CS50 for both genes (Figure 2D) with transcript reduction in these worms not significantly different to those in RPMI or those in CS50 (*fhcam2* transcript reduction in CS20: 71.3%±3.7; *fhσgst* transcript reduction in CS20: 68.3%±8.3), suggesting that growth rate may be a factor in *F. hepatica* RNAi dynamics. All trials showed a consistent trend whereby *FhCaM2* transcript knockdown was more robust than that for *FhσGST*.

To examine if these effects were similar in older juveniles, juveniles were grown for one week in CS50 prior to dsRNA exposure. Figure 3A shows how, in this case, CS50 maintained juveniles exhibited less knockdown of *fhcam2* (transcript reduction in RPMI: 96.8%±0.4; in CS50: 94.4%±0.6 – t-test: P=0.028), and no significant reduction for *fhσgst* (transcript reduction in RPMI: 91.8%±2.4; in CS50: 77.8%±8.3 – t-test: P=0.16). The transcript knockdown in this trial compared favourably with that seen if juveniles were repeatedly soaked with dsRNA over a two-week period (Figure 3A) for both *fhcam2* (transcript reduction in repeated soaks: 95.2%±0.1) and *fhσgst* (transcript reduction in repeated soaks: 70.3%±2.4). However, when comparing protein knockdown between these two regimens we found that those treated with dsRNA just once exhibited less FhCaM2 knockdown and no FhσGST knockdown (Figure 3B), whereas repeated soaking in dsRNA led to significant knockdown of both proteins (protein knockdown FhCaM2: single soak, 77%±3.5; repeated soak, 97.3%±0.4; protein knockdown FhσGST: single soak, 9.3%±16.3; repeated soak, 44.6%±10.5).

**Figure 3:**
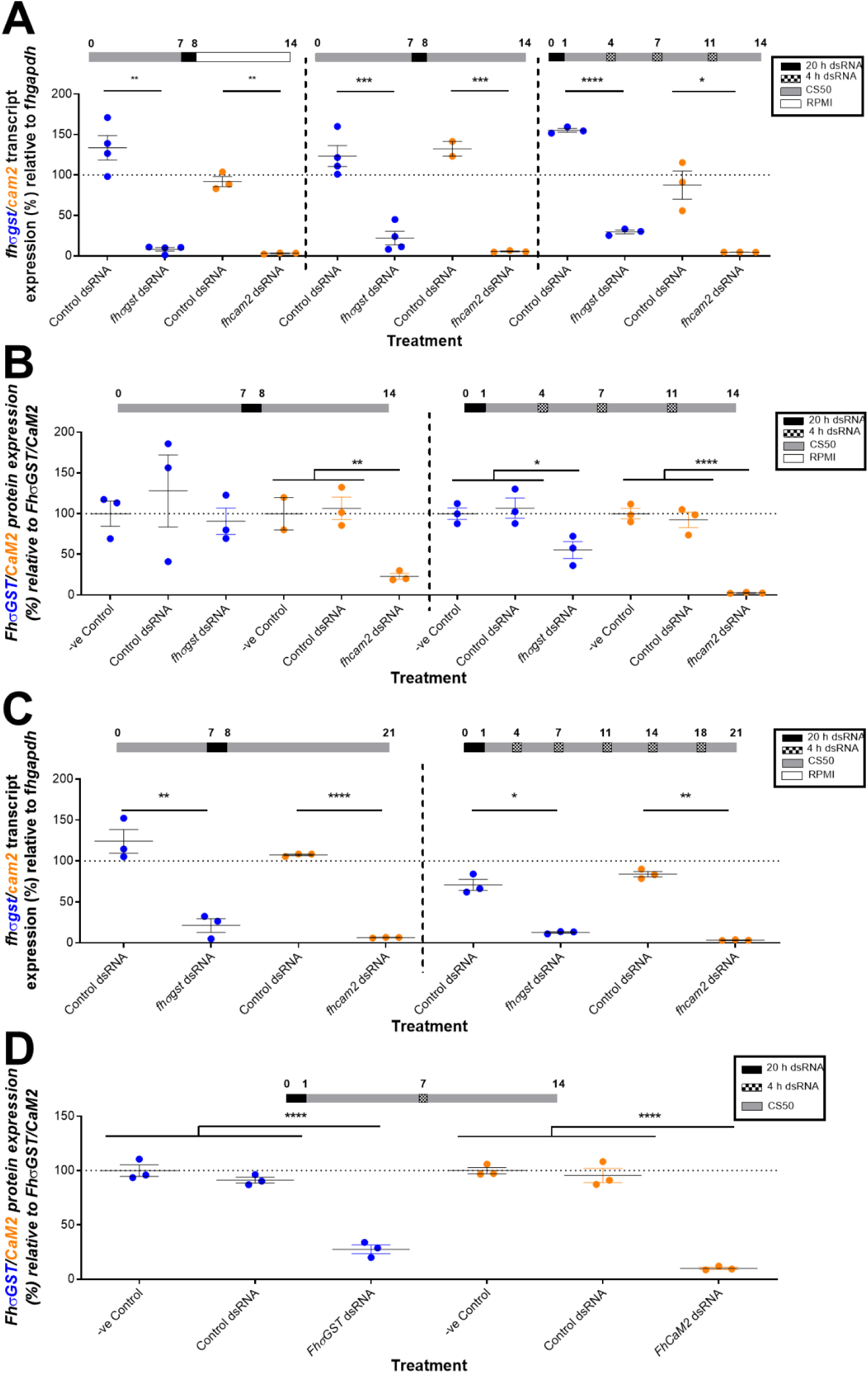
The impact of single and repeated dsRNA exposures on transcript and protein knockdown in juvenile *Fasciola hepatica* during *in* vitro growth. A – Percentage expression of *F. hepatica* sigma glutathione transferase (*fhσgst*) / *F. hepatica* calmodulin 2 (*fhcam2*) transcript relative to *F. hepatica* glyceraldehyde phosphate dehydrogenase (*fhgapdh*) following distinct dsRNA treatment protocols shows that when grown for one week prior to dsRNA exposure, the maintenance media post exposure does not alter the knockdown response. B – Over a two-week *in vitro* maintenance timecourse, the knockdown of *fhσgst* / *fhcam2* proteins relative to controls in juvenile fluke was most efficient following repeated exposures to dsRNA, with single dsRNA exposures being less effective. C - Over a three-week *in vitro* maintenance timecourse, the knockdown of *fhσgst* / *fhcam2* transcripts relative to controls in juvenile fluke was similar following single or repeated exposure to dsRNA. D – Over a three-week *in* vitro maintenance timecourse, the knockdown of FhσGST / FhCaM2 proteins relative to controls in juvenile fluke was robust following a single exposure to dsRNA after one week of growth. Treatment timelines (in days) above graphs show media that juveniles were maintained in; y axis dotted line at 100% represents mean transcript / protein expression in untreated juveniles; colours of gene/protein in y axis correspond to gene/protein expression on graphs. Unpaired t tests used to test for transcript knockdown and ANOVAs for protein knockdown *, P<0.05; **, p<0.01; ***, p<0.001; ****, p<0.0001.

When the treatment regimens were extended to run for a total of three weeks we found that there was little difference in the level of transcript knockdown (Figure 3C) for either gene (transcript reduction *fhcam2*: single soak, 93.6%±0.2; repeated soak, 96.7%±0.3; transcript reduction *fhσgst*: single soak, 78.8%±8.4; repeated soak: 87.3%±0.9), suggesting that transcript recovery was not as dramatic in more developed juveniles. Therefore, to sustain gene knockdown at least one repeated soak per week in dsRNA appears to be required. The final treatment regimen over two weeks involved a dsRNA soak immediately post-excystment, with a ‘top-up’ soak seven days later. This induced significant protein knockdown (Figure 3D) for both FhCaM2 and FhσGST (protein knockdown CaM2: 89.9%±1.1; σGST: 72.4%±4), comparable to that seen in juveniles repeatedly soaked in dsRNA (Figure 3A & B).

Our liver fluke maintenance platform allows us to maintain juveniles for many months *in vitro*, and to ensure that RNAi-mediated studies could be carried out in these worms we grew juveniles in CS50 for five months before repeatedly soaking the worms in dsRNA over a two-week period. This method resulted in both transcript and protein knockdown of FhσGST (Figure 4A & B), but only in transcript knockdown of FhCaM2 (transcript knockdown: *FhCaM2*, 94.2%±2.9; *FhσGST*, 93.9%±1.4; protein knockdown: FhCaM2, 0% (upregulation 149%±48); FhσGST, 88%±3). This is the greatest transcript knockdown, and consequently protein knockdown, of *FhσGST* seen in any of our treatment regimens. This suggests that that transcript abundance may play a role in RNAi susceptibility / dynamics as *fhσgst* expression increases over time in worms grown *in vitro*, whereas *fhcam2* decreases (Figure 4C).

**Figure 4:**
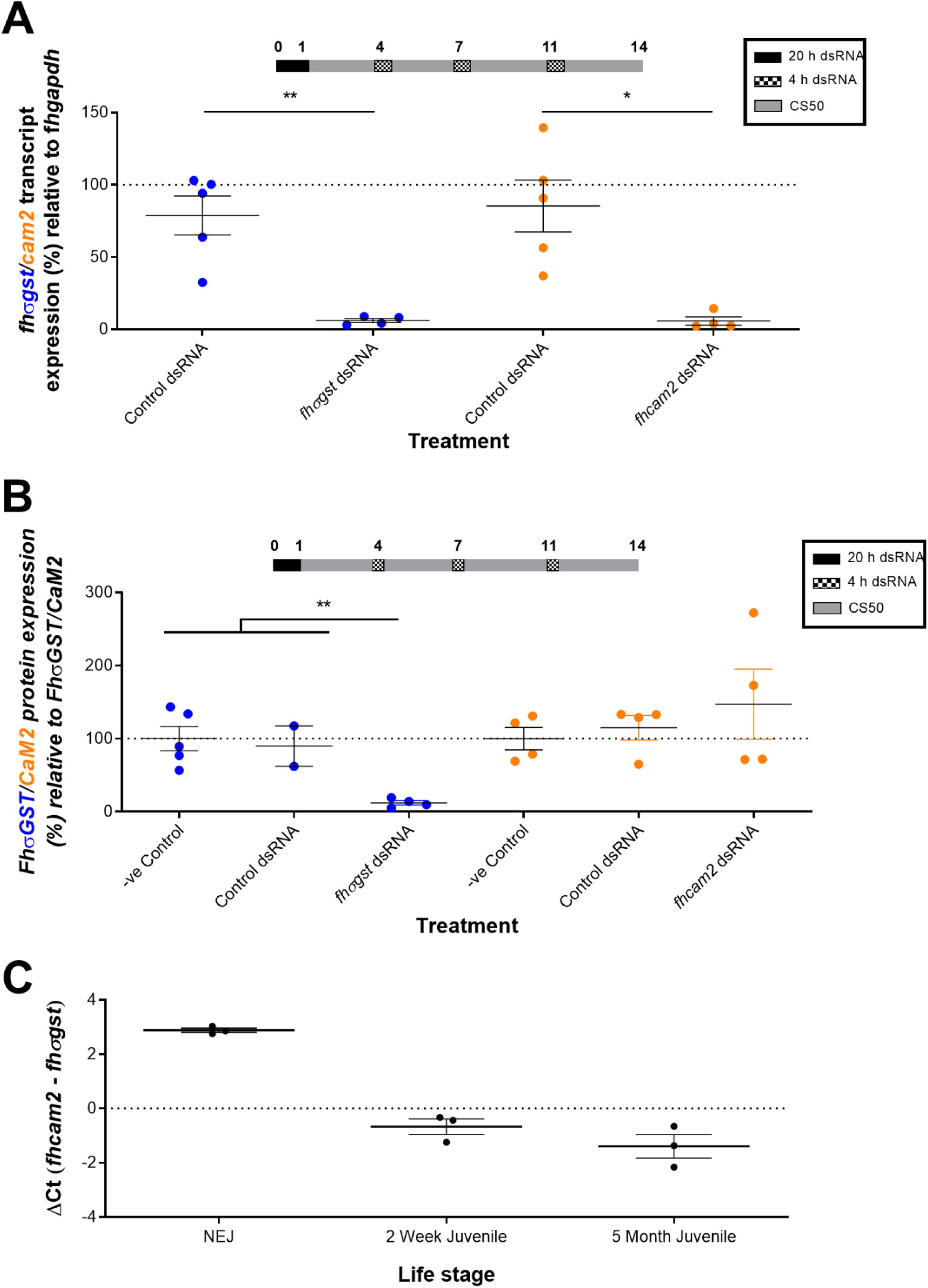
Juveniles maintained *in vitro* for five months exhibit greater protein knockdown of sigma glutathione transferase (FhσGST) than of calmodulin 2 (FhCaM2). A – Percentage expression of *fhσgst* / *fhcam2* transcript relative to *F. hepatica* glyceraldehyde phosphate dehydrogenase (*fhgapdh*) after repeated soaking in dsRNA results in knockdown of both *σgst* and *cam2* after two-weeks. B – Percentage expression of FhσGST / FhCaM2 protein relative to control protein (FhσGST / FhCaM2) shows that knockdown of σGST is greater than that of CaM2 when juveniles are repeatedly exposed to dsRNA across two-weeks. C – The relative expression (ΔΔCt) of *fhcam2* and *fhσgst* in *in vitro* juveniles shows that *cam2* is more abundant than *σgst* in early stage juveniles, but that over time *σgst* becomes more abundant than *cam2*. Treatment timelines (in days) above graphs show media juveniles were maintained in; y axis dotted line at 100% represents mean transcript/protein expression in untreated juveniles; colours of gene in y axis correspond to gene expression on graphs. Unpaired t tests used to test for transcript knockdown and ANOVAs for protein knockdown *, P<0.05; **, p<0.01.

Together, these data highlight an increased robustness of RNAi-mediated gene silencing in *F. hepatica* juveniles at different stages of development using a repeated soaking methodology. Our improved methodology of repeated dsRNA soaks (24 h/soak) over an extended period has now allowed us to efficiently silence genes in both stem cells (*fhmap3k1*) and muscle tissue (*fhparamyo*) of juvenile *F. hepatica* (Supplementary Figure 4).

## 4. Discussion

Due to the threat that *F. hepatica* poses to global food security and to human health, it is critical that novel targets for control are identified. RNAi could aid in the identification and validation of these targets and we have shown here that all life stages of *F. hepatica* express the necessary molecular machinery to support RNAi. Interestingly, it appears that liver fluke, along with other parasitic flatworms, have undergone expansion of the RNAi effector machinery. Our results strengthen the idea that the piwi-clade AGOs have been lost in the parasitic flatworms, only to be replaced by a parasitic flatworm specific clade (Skinner et al. 2014), with an as yet unknown function. It is possible that this novel pathway acts on parasitic flatworm transposons (DeMarco et al., 2004; Jacinto et al., 2011; Koziol et al., 2015), as the piRNA, endo si-RNAs and clade I AGOs do in germ line and somatic cells in other eukaryotes (Vagin et al., 2006; Houwing et al., 2007; Chung et al., 2008; Ghildiyal et al., 2008; Kawamura et al., 2008). Given the difference in the number of repeat sequences between the UK and Oregon *F. hepatica* genomes (30.6% and 55.3%) (McNulty *et al*., 2017) it may be that the regulation of transposable elements does not occur in the parasitic flatworms, hence the absence of PIWI AGOs in *F. hepatica*.

Further evidence to support the idea that these parasitic flatworm AGOs have replaced the Piwi proteins comes from *Schistosoma* sp. as *ago2* is highly expressed in *S. mansoni* (Cogswell et al., 2011) and has ubiquitous expression in *S. japonicum* where it is thought to be expressed in both germline and somatic cells (Cai et al., 2012). Given that all of the *F. hepatica* AGOs, including those specific to parasitic flatworms, are expressed in all life stages, it is possible that they exhibit ubiquitous expression, as reported in schistosomes. Our analysis shows that other RNAi effectors have undergone expansion in the parasitic flatworms, such as FMR1s which may form part of the RISC. These predicted FMR-like proteins are strikingly divergent from orthologues in other eukaryotes, forming two distinct groups, with *F. hepatica* possessing four genes in total. All four f*hfmr*-like genes were expressed in all *F. hepatica* life stages, suggesting involvement in baseline processes consistent throughout the life cycle; localisation studies would help inform any associated tissue specificities.

The expansion of the RNAi effectors in the parasitic flatworms suggests that the RISC associated with piRNAs in other eukaryotes has been replaced by a novel RISC in the parasitic flatworms. As piRNAs have not been observed in the parasitic flatworms it is possible that they possess a different, and unique, subset of noncoding RNAs. Further evidence for a separate noncoding RNA pathway is provided by the expansion of DCR and ERI clades in parasitic flatworms. Both cestodes and trematodes possess a novel *dcr* gene as well as a distinct *eri*-*2* gene. Flukes also appear to express an additional *eri*-*1* that seems to have been lost in cestodes. This additional DCR could act on ncRNA precursors prior to RISC incorporation in a similar way to how DCR processes endo-siRNAs found in both germ and somatic cells of mammals and flies (Czech 2008, Tam 2008, Ghildiyal 2008, Song 2011), whilst novel ERIs may act as inhibitors of this pathway.

As our *in silico* analysis suggested that all life stages of the liver fluke have a functioning RNAi pathway we wanted to further explore the utility of RNAi in different *F. hepatica* developmental life stages. As our culture platform developments (McCusker et al. 2016) include changes to the base medium used to maintain juveniles during *in vitro* trials we wanted to establish if this affected RNAi efficiency. We found that *in vitro* growth in CS50 results in transcript recovery over the first seven days post excystment, but this has less of an effect once juveniles are more than seven days old. Transcript recovery has not been previously noted in RNA dynamics studies in *F. hepatica* (Dell’Oca et al. 2014; McVeigh et al. 2014), likely due to the fact that the juveniles used in those studies had not been maintained in growth-promoting media. This observation of transcript recovery in *F. hepatica* differs from that seen in *Schistosoma* sp. where *in vitro* RNAi is persistent for many weeks (Correnti et al., 2005; Krautz-Peterson et al., 2010), highlighting both the potential challenges in optimizing RNAi under conditions where growth and development are not occurring as well as the differences in *Fasciola* and *Schistosoma*, two species often compared during RNAi phenotypic readouts. Changes in transcriptional activity in the first week post-excystment may contribute to the transcript recovery observed and work in mammalian cell lines has highlighted that such gene expression dynamics may associate with diminished RNAi susceptibility (Larsson et al., 2010). This period of rapid gene expression change following excystment correlates with the increased metabolic activity that occurs as NEJs transition from a sedentary life stage to an actively developing, migrating and feeding life stage.

The observation that the ability of transient dsRNA exposure to trigger RNAi in juvenile fluke was not altered when CS50 was used to reduce the amount of residual dsRNA in the bathing medium post dsRNA treatment, indicates that the initial dsRNA exposure is the key trigger for gene silencing. Therefore, subsequent gene silencing dynamics are a result of the continued processing and internal transport of RNAi triggers acquired by the fluke during the transient dsRNA exposure phase. Also, changes in the efficiency of gene knockdown in fluke maintained in different media (RPMI, CS20 or CS50) supports our proposed association between transcript recovery and fluke growth over the first week post-excystment.

It was also noteworthy that the expression of many RNAi effectors, such *fhago1* and *fhago2*, increased after one week of growth in CS50, compared to juveniles maintained in RPMI (Supplementary Figure 5). *fhago2*, which doubled in expression, is known to be crucial for siRNA-directed gene silencing in *D. melanogaster* (Okamura et al., 2004), suggesting that it is crucial in exogenous dsRNA / siRNA gene silencing. Given that juveniles grown in CS50 for one week prior to dsRNA exposure did not exhibit differences in RNAi efficiency between different media, it appears that RNAi effector expression levels have little impact on RNAi efficiency *in vitro*. Our results suggest that *in vitro* fluke growth is the key factor that alters RNAi dynamics in early stage juveniles.

From our experiments we concluded that repeatedly soaking juveniles in dsRNA is necessary to ensure sustained target transcript knockdown. Repeated dsRNA exposure facilitated the most efficient protein knockdown, most likely because this reduced the opportunity for the recovery of target expression. However, as noted in previous studies, it can take over a week to elicit a reasonable level of protein knockdown for some genes in *F. hepatica* (McVeigh et al. 2014; McCammick et al. 2016). Therefore, we recommend that to achieve RNAi in growing *F. hepatica* juveniles repeated dsRNA exposures are necessary to ensure the robust knockdown of transcript and protein across periods in excess of a week. We confirmed this by testing repeated 24 h dsRNA soaks over the course of a month for two genes expressed in stem cells (*fhmap3k1*) and muscle tissue (*fhparamyo*), with extensive transcript knockdown in both cases, confirming the robustness of our functional genomics platform in this crucial life stage.

We also observed differences in the level of knockdown (both transcript and protein) between target genes as *fhcam2* exhibited consistently higher knockdown than *fhσgst* in the early stage juveniles. However, in juveniles grown for five months prior to RNAi trials we found that FhσGST protein was silenced more effectively than FhCaM2. These differences appear to correlate with the relative abundance of each transcript as in the early stage juveniles *fhcam2* is more abundant than *fhσgst*, whereas in the later stage juveniles (five months) the reverse is true. Work in mammalian cell lines has observed correlations between transcript abundance and RNAi susceptibility with higher expressed transcripts being more amenable to RNAi mediated knockdown (Hong et al., 2014).

We have also carried out the first successful RNAi in adult *F. hepatica* using two different methodologies (Supplementary File 2). Both nanoinjection and soaking of adults appeared to elicit knockdown of *FhCatL, FhσGST* and *FhCaM2* (Supplementary Figure 6A & B). However, although these approaches appeared to elicit robust transcript knockdown over 48 h there was no reduction in protein levels observed (Supplementary Figure 6C). This lack of protein knockdown may relate to transcript turnover rates being affected by deteriorating health when worms are removed from the host. This suggests that the chief barrier to a robust RNAi platform in adult *F. hepatica* is the lack of a reliable *in vitro* maintenance system.

The results presented here highlight the fluid dynamics of RNAi in a growing parasitic flatworm for the first time. As such we recommend the use of a ‘repeated dsRNA soaking’ methodology to ensure robust RNAi in *F. hepatica* juveniles and facilitate phenotypic studies. The confirmation of a functional RNAi pathway in all life stages of the liver fluke means that this functional genomics tool can be utilised to ascertain gene function across all key life stages although, a study by Beesley et al. (2016) demonstrated the genetic diversity of wild populations of *F. hepatica* within the UK, another factor that could have implications for RNAi susceptibility. Here we have demonstrated a robust platform for gene silencing in the highly pathogenic juvenile stage liver fluke which will facilitate the exploitation of the new genomic/transcriptomic datasets to advance studies on liver fluke biology.

## 5. Acknowledgements

Funding for this study was provided by the Biotechnology and Biological Sciences Research Council and Merial Ltd. (BB/K009583/1), the National Centre for the 3Rs (NC/N001486/1), the John Glover Memorial Fund and the Department for the Economy of Northern Ireland. Thanks to David Timson and Peter Brophy for the original preparation of antisera for FhCaM2 and FhσGST used in main manuscript. Thanks to John P. Dalton for original preparation of antisera for FhCatL used in supplementary data. Thanks also to Cieran Donald at ABP Meats, Lurgan for facilitating the collection of adult liver fluke samples.

## Declarations of interest

none

## Supplementary Files

**Supplementary Figure 1: Maximum-likelihood trees of putative *F. hepatica* RNAi effectors alongside the RNAi effectors of *Schistosoma* sp., *Caenorhabditis elegans* and *Homo sapiens*.** (A) Drosha, (B) DGCR8, (C) TRBP, (D) TSN-1, (E) SID-1, (F) Exportin-1, (G) Exportin-2, (H) Exportin-5, (I) SMG-2, (J) SMG-4, (K) SMG-5, (L) SMG-6, (M) FMR1s, (N) ERIs. Trees A-L are JTT models with M and N WAG models.

**Supplementary Figure 2: Sequence alignments of *Schistosoma* sp., *Caenorhabditis elegans, Homo sapiens* and putative *F. hepatica* RNAi effectors** – (A) Alignment of putative F. hepatica Drosha against known homologues with endonuclease regions (grey) and catalytic residues (black) marked. (B) Alignment of putative F. hepatica DGCR8 against known homologues with dimerization domain (red box), dsRNA binding domains (blue and purple boxes), dimerization interacting residues (grey) and nucleotide binding residues (pink) highlighted. (C) Alignment of putative F. hepatica Argonautes against known homologues with midpoint domain (light blue box), PIWI domain (purple box), 3’ end binding residues (yellow), 5’ end residues (orange), m7g Cap binding residues (red), Mg2+ interacting residues (purple) and ‘DDH’ catalytic residues (black) highlighted. (D) Alignment of putative F. hepatica Dicers against known homologues with RNase III domains (blue boxes), dsRNA binding domain (pink box) and catalytic residues (black). (E) Alignment of putative F. hepatica ERIs against known homologues with possible Dicer interacting residues (black) highlighted. (F) Alignment of putative F. hepatica TRBP against known homologues with ‘DEDHD’ catalytic residues (black) highlighted. Across all alignments green coils and blue arrows mark alpha helices and beta strands based on published crystal structures.

**Supplementary Figure 3: Agarose genes showing expression of putative RNAi effectors in different *F. hepatica* life stages.** (A) *fhgapdh*, (B) *fhdgcr8*, (C) *fhdrosha*, (D) *fhdcr-1*, (E) *fhdcr-2.1*, (F) *fhtrbp*, (G) *fhdcr-2.2*, (H) *fhago1*, (I) *fhago2*, (J) *fhtsn-1.1*, (K) *fhago3*, (L) *fhfmr1.1*, (M) *fhfmr1.2*, (N) *fhfmr1.4*, (O) *fhfmr1.3*, (P) *fhexportin-1*, (Q) *fhexportin-2*, (R) *fhsid-1*, (S) *fhexportin-5*, (T) *fheri-1*, (U) *fheri-2*, (V) *fheri-3.2*, (W) *fheri-3.1*, (X) *fhsmg2*, (Y) *fhsmg4*, (Z) *fhsmg6*, (AA), *fhsmg5*. For each gene running order from left to right: 100bp ladder, unembryonated eggs, developed eggs, NEJs, 3-week-old juveniles, adult, no RT control, negative template control.

**Supplementary Figure 4: RNAi-mediated knockdown using repeated soaking methodology of *fhparamyo* (paramyosin) and *fhmap3k1* (Mitogen-activated protein 3k1) in juvenile Fas*ciola hepatica*.** Percentage expression of (A) *fhparamyo* and (B) *fhmap3k1* transcript relative to *F. hepatica* glyceraldehyde phosphate dehydrogenase (*fhgapdh*) after repeated 24 h dsRNA soaks across (A) four or (B) three weeks. Treatment timelines (in weeks) above graphs show media juveniles were maintained in; y axis dotted line at 100% represents mean transcript expression in untreated juveniles. Unpaired t tests used to test for transcript knockdown *, P<0.05.

**Supplementary Figure 5: *fhago1*/*fhago2* expression is greater in *F. hepatica* juveniles maintained in growth-inducing media –** Percentage expression of *fhago1* / *fhago2* and *fhdgcr8* in juveniles grown for one week in CS50 relative to juveniles maintained in RPMI shows an increase in *fhago1* and *fhago2* expression, but not *fhdgcr8*. Y axis dotted line at 100% represents expression in juveniles maintained in RPMI.

**Supplementary File 6: Adult *Fasciola hepatica* cathepsin L (FhCatL) is susceptible to RNAi via either soaking in double stranded RNA (dsRNA) or dsRNA nanoinjection.** A – Percentage expression of *fhcatl* / calmodulin 2 (*fhcam2*) transcript relative to *F. hepatica* glyceraldehyde phosphate dehydrogenase (*fhgapdh*) after 18 h soak in dsRNA, followed by 30 h maintenance in CS20 results in CatL knockdown but no significant *fhcam2* knockdown. B – Percentage expression of *fhcatl* transcript relative to *fhgapdh* after nanojection with dsRNA, followed by 18 h incubation in RPMI and a further 30 h maintenance in CS20 showing knockdown of CatL. C - Percentage expression of FhCatL protein relative to control after nanojection with dsRNA, followed by 18 h incubation in RPMI and a further 30 h maintenance in CS20 shows no knockdown of FhCatL. y axis dotted line at 100% represents mean transcript/protein expression in untreated juveniles; colours of gene/protein in y axis correspond to gene/protein expression on graphs. Unpaired t tests used to test for transcript knockdown; **, p<0.01; ****, p<0.0001.

**Supplementary Table 1: Primers used in PCRs of putative *F. hepatica* RNAi effectors.**

**Supplementary Table 2: Primers used in dsRNA synthesis and qPCRs.**

**Supplementary Table 3: Putative *F. hepatica* RNAi effectors.**

**Supplementary File 1: RNAi effector sequences used to generate Hidden Markov Model (HMM) queries for HMMsearch of *F. hepatica* predicted proteome.**

**Supplementary File 2: Methodology for RNA interference in adult *F. hepatica*.**

## Notes

### Competing Interest Statement

The authors have declared no competing interest.

