## Supplementary Data for "RNA interference dynamics in juvenile *Fasciola hepatica* are altered during *in vitro* growth and development"

#### Supplementary Figure 1

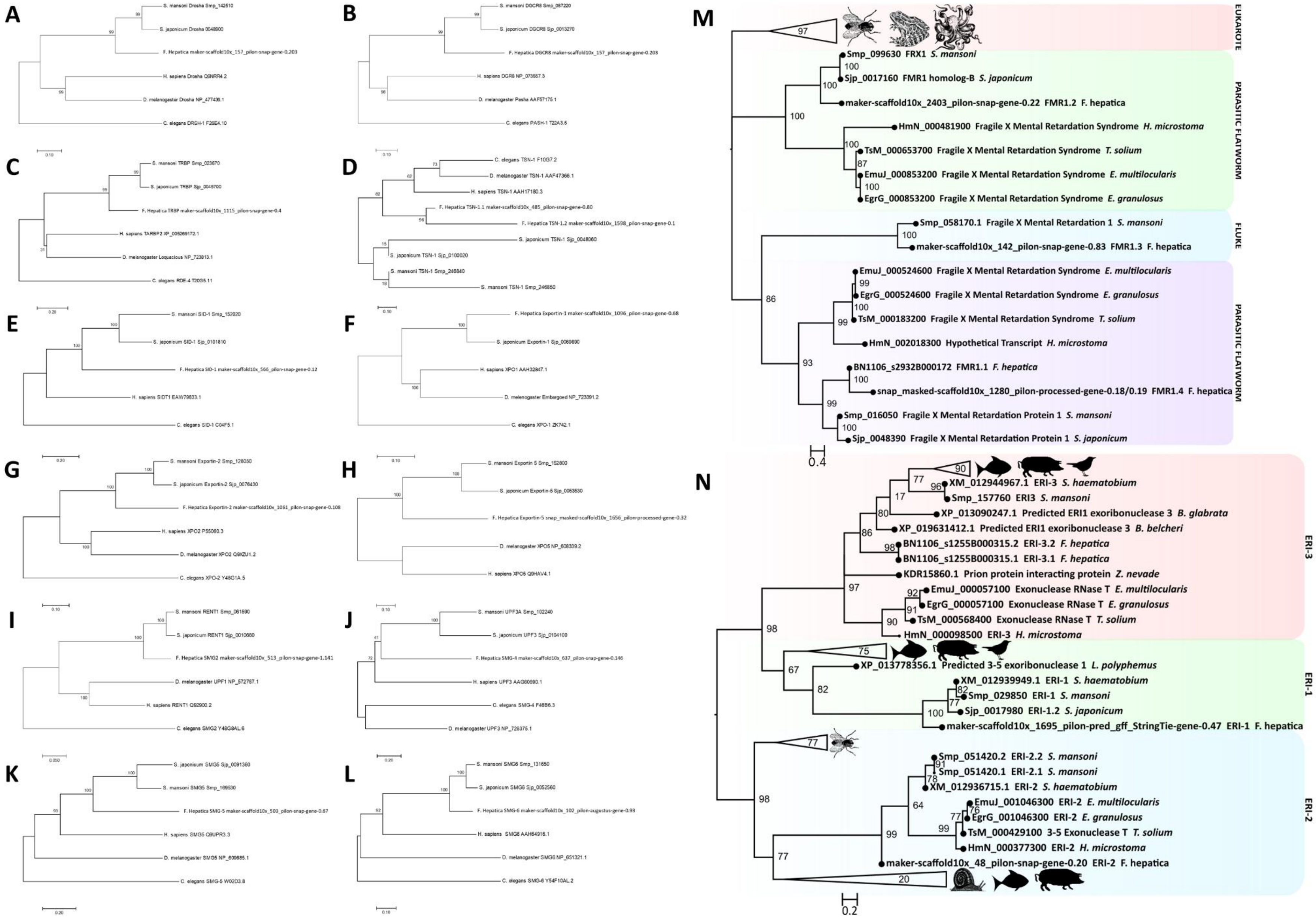

#### Supplementary Figure 2

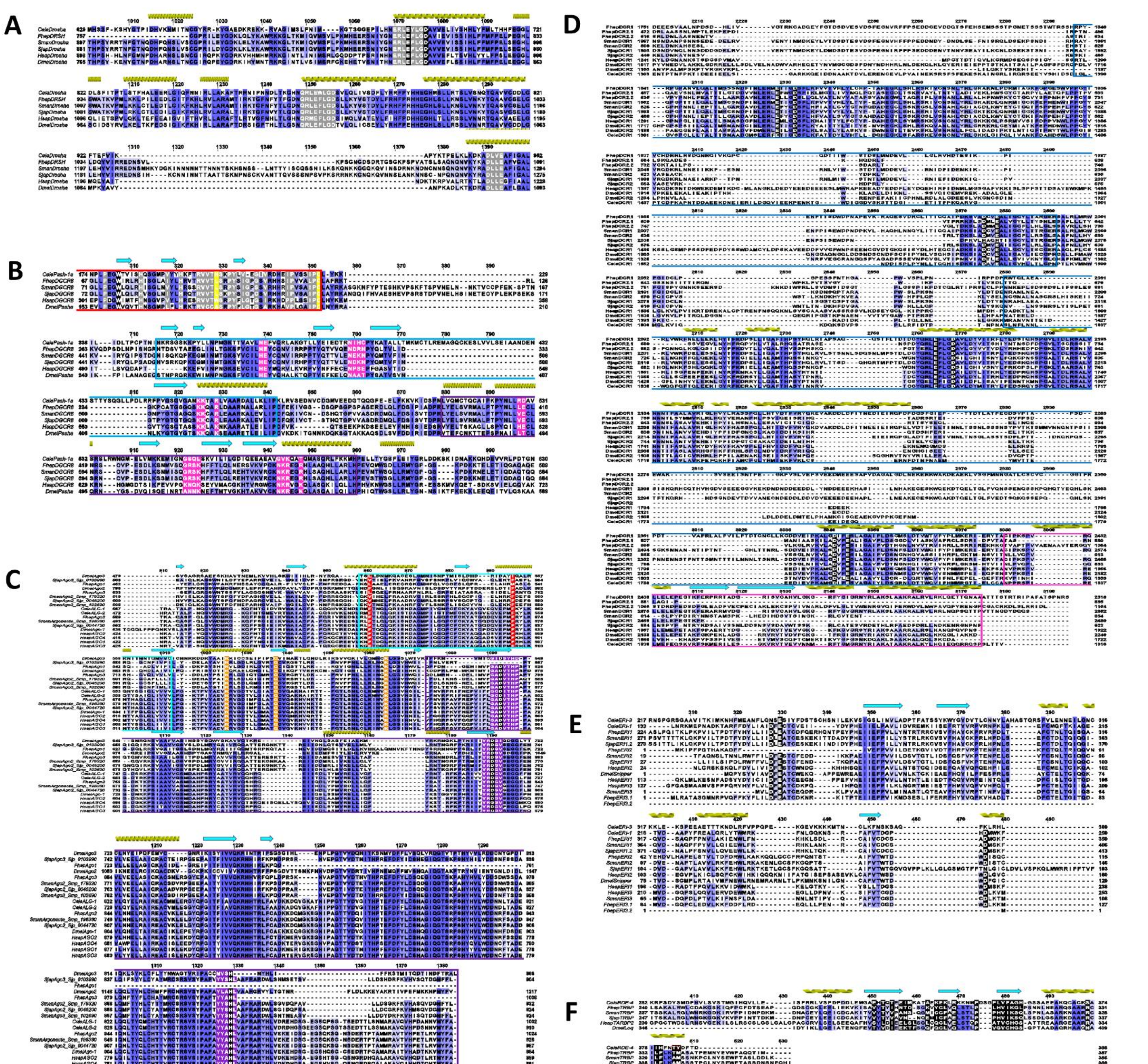

**Supplementary Figure 3**

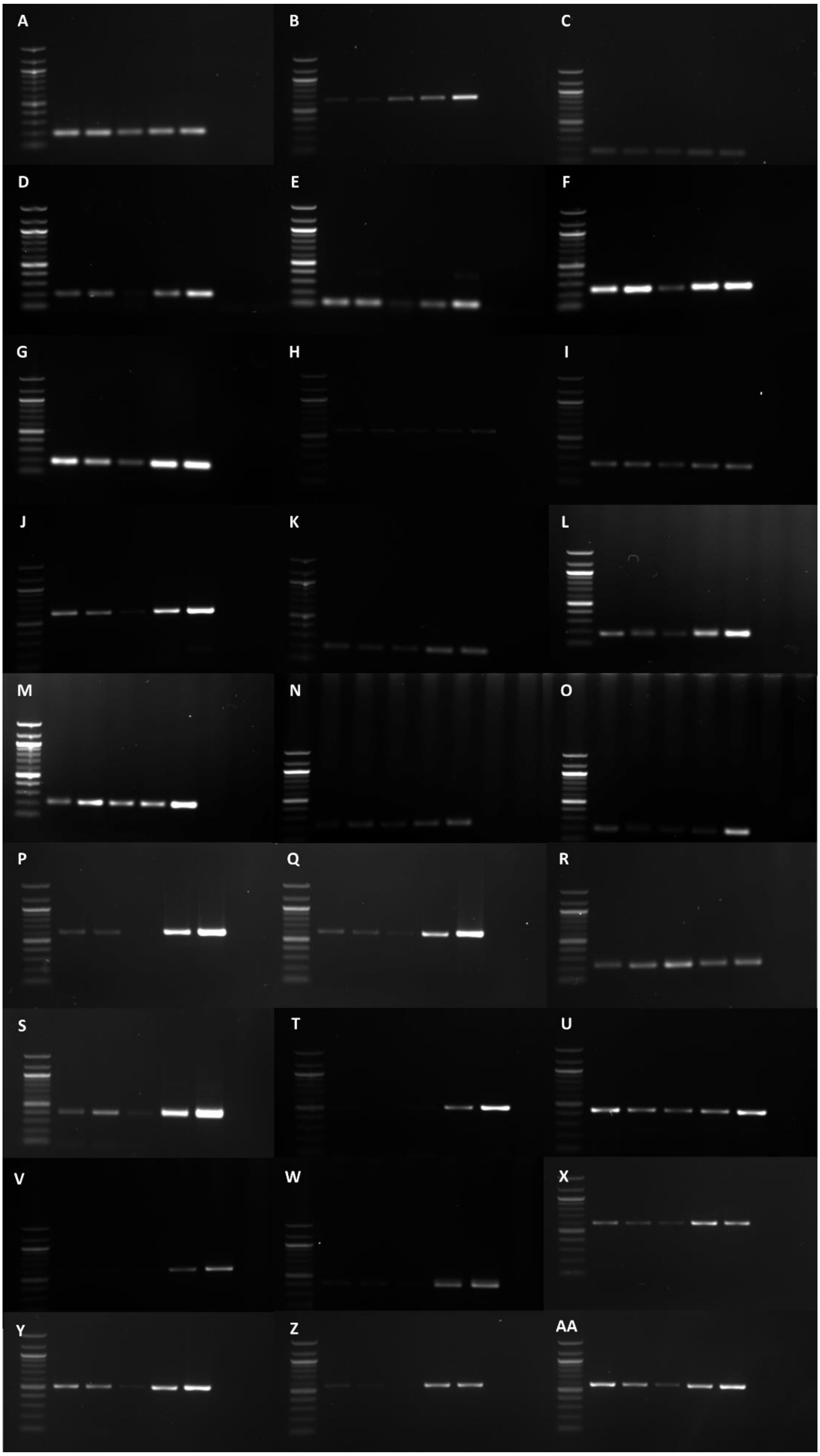

**Supplementary Figure 4**

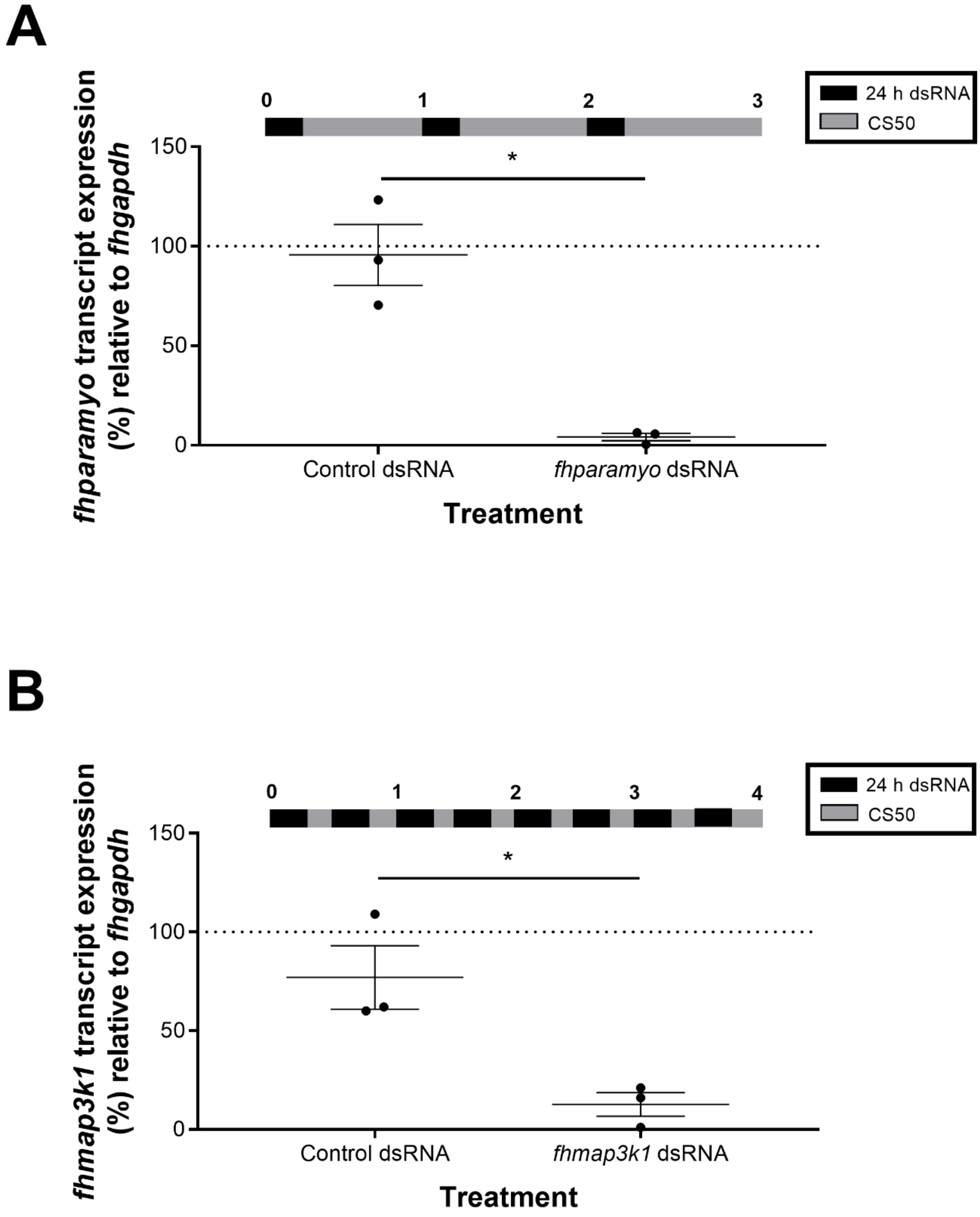

**Supplementary Figure 5**

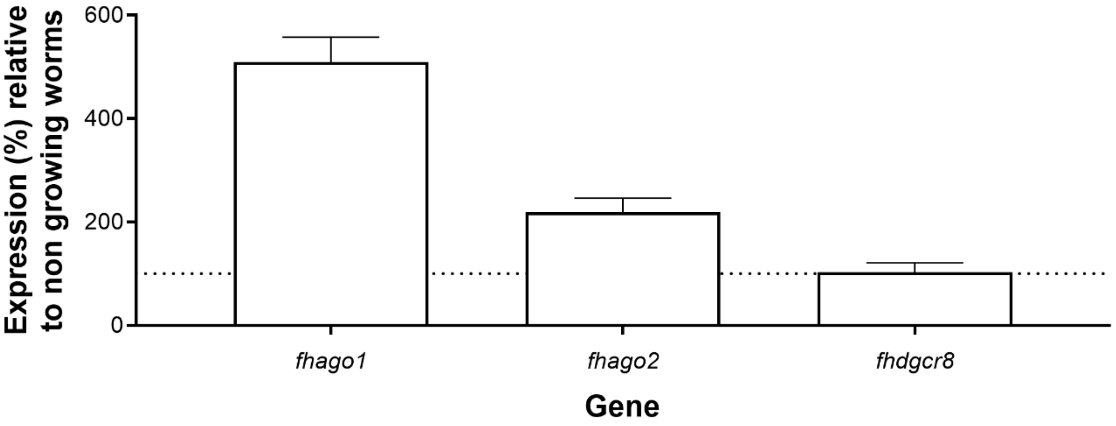

**Supplementary Figure 6**

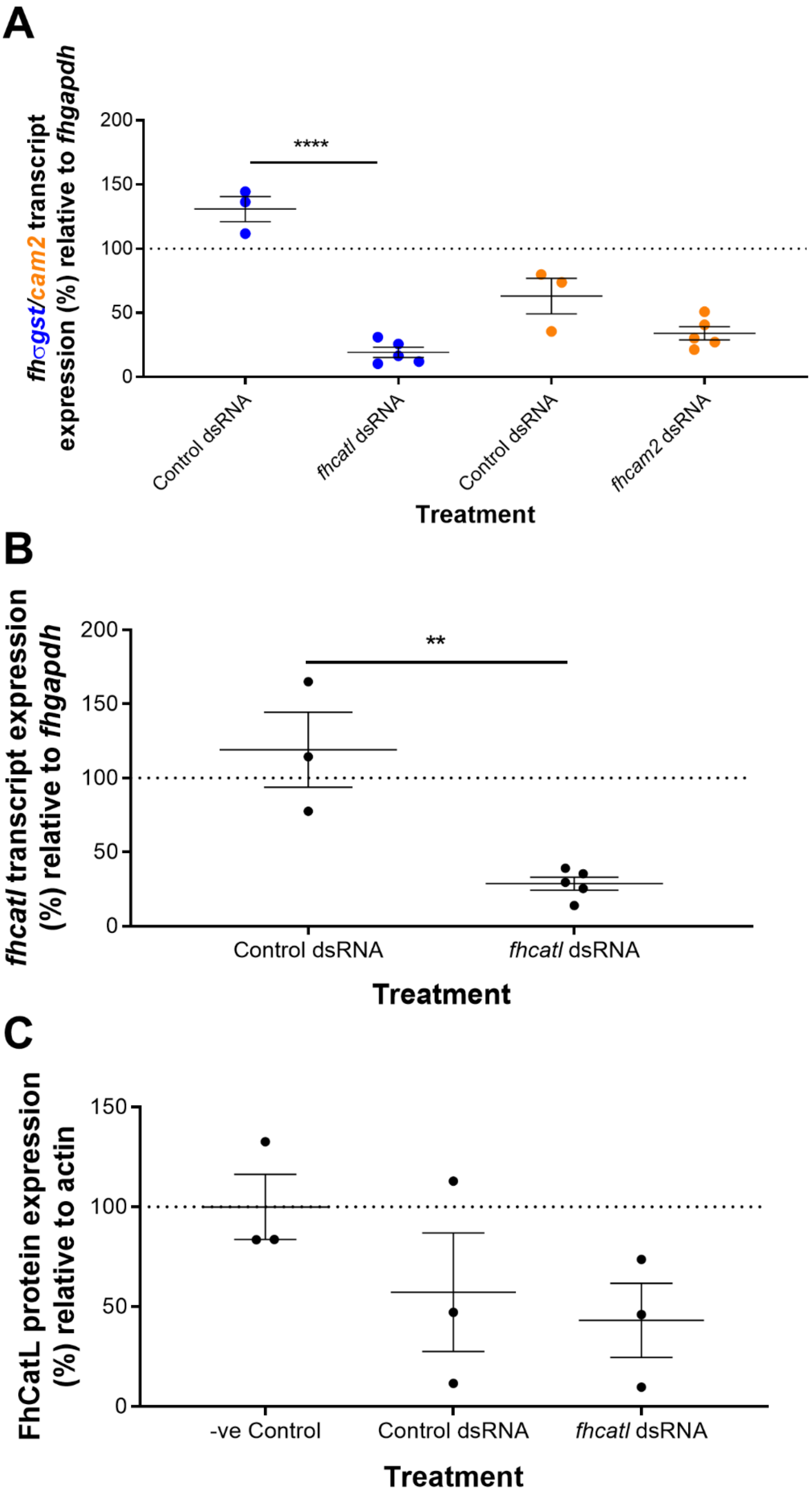

#### Supplementary Table 1

| Designation | *F. hepatica* ID (Predicted Protein Dataset) | Forward Primer | Reverse Primer | Product Size |
|---|---|---|---|---|
| *drosha* | maker-scaffold10x_157_pilon-snap-gene-0.203 | GCAGGCGATTGTGCAGAATC | GCGATCACTGTCACCATTGC | 745 |
| *dgcr8* | maker-scaffold10x_503_pilon-snap-gene-0.75 | GGCCATACTCTATCGGTCCA | ATCAGCGGCTGTTAGCAATC | 200 |
| *dcr-1* | maker-scaffold10x_432_pilon-snap-gene-0.93 | TTACCGACCGGAAGATGAAC | TCTTCCGTTAGTGGGTGTCC | 204 |
| *dcr-2.1* | maker-scaffold10x_1080_pilon-augustus-gene-0.2 | ACGCAGTTTTTACCGCCAAC | ACCAACGCTTCCAGCATATC | 119 |
| *dcr-2.2* | maker-scaffold10x_1080_pilon-snap-gene-0.12 | CCAAGTGACTGCTGGTTTGC | GGGTTGGAACGCCTCACTTA | 250 |
| *trbp* | maker-scaffold10x_1115_pilon-snap-gene-0.4 | TTTGCTTGGTTCCGATTTTC | TGCAGTGGTACTCGTGCATT | 197 |
| *ago-1* | maker-scaffold10x_919_pilon-snap-gene-0.104 | GAACTAGCGCATCGAATGGC | CCAGCCAATTCCTCGAGGAG | 599 |
| *ago-2* | maker-scaffold10x_145_pilon-augustus-gene-2.246 | AATCAGGCAATTCCGATGAG | TGAGAATTTCGCGACGGTGA | 249 |
| *ago-3* | maker-scaffold10x_979_pilon-snap-gene-0.4 | AATGACTTGCGCCGAATACG | TGTTTTTGAGCAGCCCTGTC | 578 |
| *tsn-1* | maker-scaffold10x_485_pilon-snap-gene-0.80 | GTTGTCGCATCCGGTATTCT | TGAACCCGTAAAGGGTGAAG | 200 |
| | maker-scaffold10x_1598_pilon-snap-gene-0.1 | - | - | - |
| *fmr1* | maker-scaffold10x_889_pilon-snap-gene-0.31 | CCGCGATTCGGTAGAGAAG | ATTCATTCCTCGACCACCAC | 190 |
| | maker-scaffold10x_2403_pilon-snap-gene-0.22 | CATACCGTGCTGTTCCAACC | GCTGATGCGGTTGTTGATAA | 211 |
| | maker-scaffold10x_142_pilon-snap-gene-0.83 | TACCAGCTGTGTCCTTGTCG | CTTCAAGAGATCCGGACCAG | 216 |
| | snap_masked-scaffold10x_1280_pilon-processed-gene-0.18/0.19 | GTCAGACGGGCAGTAATGG | TCGCTGGCTCGACAACTATC | 202 |
| *sid-1* | maker-scaffold10x_566_pilon-snap-gene-0.12 | CGGATTTGGGAAGTAATGTGCC | GTACGCCTCGAGTCTGTGTC | 636 |
| *exportin-1* | maker-scaffold10x_1096_pilon-snap-gene-0.68 | GCCCACCGACAAGTTCAAAC | ACACCAGCTTGACGTCGTAG | 570 |
| *exportin-2* | maker-scaffold10x_1061_pilon-snap-gene-0.108 | TTCGTCGTATTCCTCGCTCG | TTGTTTGGCTCGTCCTCCTC | 551 |
| *exportin-5* | snap_masked-scaffold10x_1656_pilon-processed-gene-0.32 | TGCCCTGGTGCCTTTATTAC | ACACGAAACCATTGCACGTA | 209 |
| *eri1* | maker-scaffold10x_1695_pilon-pred_gff_StringTie-gene-0.47 | TTGAACCAGTTGGGCTCTCC | ACAGAATGCAGTCAGGTCCG | 501 |
| *eri2* | maker-scaffold10x_48_pilon-snap-gene-0.20 | ACCGCTTTGTGAGGCCTAC | TGTTCTTCGCGTCGTCCATG | 441 |
| *eri3** | BN1106_s1255B000315.1 | TGAGAGCCTCATCTTCGAACG | AAGCAATACAATGTTCGAGGATG | 591 |
| | BN1106_s1255B000315.2 | GGATGTCCTGAAGGTGTGTTTTC | TCCAGTGATTCTCGGGATAGC | 410 |
| *smg-2* | maker-scaffold10x_513_pilon-snap-gene-1.141 | GTCCCAGCCCGGATTATCTG | CCATGACTCCTCCGTGATCG | 588 |
| *smg-4* | maker-scaffold10x_637_pilon-snap-gene-0.146 | CGAAACCGTCAACACCAACC | ATTGCCGTAGCCTCCATGTC | 503 |
| *smg-5* | maker-scaffold10x_503_pilon-snap-gene-0.67 | CGGTCTGAACGCCCTCTATC | TGCGTCCATATCAGGGTTCG | 562 |
| *smg-6* | maker-scaffold10x_102_pilon-augustus-gene-0.93 | GCTCGAGGGTATAGCCACAC | GGCAAACGTGACCAGTTGAC | 573 |

*These predicted genes are absent from new genome assembly, however PCR data have confirmed their presence and expression in juvenile and adult worms

#### Supplementary Table 2

| Designation | ID | *NCBI/Wormbase Parasite ID* | Forward dsRNA Primer* | Reverse dsRNA Primer* | Forward qPCR Primer | Reverse qPCR Primer |
|---|---|---|---|---|---|---|
| ***fhcam2*** | Calmodulin 2 | AM412547/maker-scaffold10x_520_pilon-snap-gene-1.36 | AGATGCCACGGAACTGAAAG | AAATGTACCCGTCGCCATTA | ATCGCGGCAATCTTGATAAT | TTGCGCAGAAGAGTCAAGAA |
| ***fhσgst*** | σ Glutathione s-transferase | DQ974116/maker-scaffold10x_1043_pilon-snap-gene-0.18 | AATTCGCCTTCTGCTCACTTGC | TACTCTTCGTCTGTTTCACC | AATTCGCCTTCTGCTCACTTGC | TCTTCACACTCACCAATGATACG |
| ***fhparamyo*** | Paramyosin | augustus_masked-scaffold10x_377_pilon-processed-gene-0.188 | CTCTGGCATCTGCACGTAAA | TTCCTCGATCTGCTGTTGTG | TTGAAATCCGCGAAATTACC | ATCGACGTGATTCACCATCA |
| ***fhmap3k1*** | Mitogen Activated Protein 3k1 | maker-scaffold10x_208_pilon-snap-gene-0.24 | AGCCCCACACAATACCAGAG | AATTGGTGGACTGGGTTGG | AGCCCCACACAATACCAGAG | TGTACCTGCTAAGCGTCGTG |
| ***fhcatl*** | Cathepsin L | See McVeigh et al. 2014 | TKRTTATGTGACGGAGGTGA | GCCBKRTAHGGRTAAK | TKRTTATGTGACGGAGGTGA | GTATAGAAGCCAGTCACTTTGGC |
| ***fhgapdh*** | GAPDH | AY005475/maker-scaffold10x_2706_pilon-snap-gene-0.15 | | | GGCTGTGGGCAAAGTCATTC | AGATCCACGACGGAAACATCA |
| ***neo*** | Bacterial Neomycin | U55762 | GGTGGAGAGGCTATTCGGCT | CCTTCCCGGTTCAGTGACAA | | |

*****dsRNA constructs were amplified with primers labelled at the 5' end with T7 RNA polymerase promoter sequence (taatacgactcactatagggt) to make T7 labelled templates in each direction

#### Supplementary Table 3

| Name | S. mansoni ID | F. hepatica ID | Length (aa) | % Identity with *S. mansoni* | Domains/Pfam IDs | Signature Motifs Identified | |
|---|---|---|---|---|---|---|---|
| **Drosha** | Smp_142510 (1531aa) | maker-scaffold10x_157_pilon-snap-gene-0.203 | 1640 | 45.00 | Ribonuclease III domain (IPR000999)<br>Double-stranded RNA-binding domain (IPR014720) | **Endonuclease 1**<br>ERLEFLGD | **Endonuclease 2**<br>QRLEFLGD… DLLE |
| **DGCR8** | Smp_087220 (761aa) | maker-scaffold10x_503_pilon-snap-gene-0.75 | 457 | 37.60 | Double-stranded RNA-binding domain (IPR014720) | | |
| **Dicer-1** | Smp_169750 (2646aa) | maker-scaffold10x_432_pilon-snap-gene-0.93 | 2422 | 46.10 | P-loop containing nucleoside triphosphate hydrolase (IPR027417)<br>Dicer dimerisation domain (IPR005034)<br>PAZ domain (IPR003100)<br>Ribonuclease III domain (IPR000999)<br>Double-stranded RNA-binding domain (IPR014720) | **3' End Pocket**<br>SPFPTSTY…NFAAYYMSKY…IPELC | **5' End Pocket**<br>RGRPCYRPEDEPLGFGLLTTKPLHHIPTFPIFSRSGEETVRF…TVMRLSLIVPRYVN |
| **Dicer-2** | Smp_033600 (954aa) | maker-scaffold10x_1080_pilon-augustus-gene-0.2 | 601 | 30.83 | Ribonuclease III domain (IPR000999) | **3' End Pocket**<br>DPVPTKLS…SYVDYFSIRY…LADTC | **3' End Pocket**<br>RHHEDFDPGKLTDSRSNIVSNNSLASAVVEHKIHPY(2088)…GACRKDLSLRRID |
| | | maker-scaffold10x_1080_pilon-snap-gene-0.12 | 1091 | 41.50 | Ribonuclease III domain (IPR000999) | **3' End Pocket**<br>DPVPTKLS…SYVDYFSIRY…LADTC | **3' End Pocket**<br>RHHEDFDPGKLTDSRSNIVSNNSLASAVVEHKIHPY(2088)…GACRKDLSLRRID |
| **TRBP** | Smp_023670 (356aa) | maker-scaffold10x_1115_pilon-snap-gene-0.4 | 357 | 56.10 | Double-stranded RNA-binding domain (IPR014720), | | |
| **Argonaute-1** | Smp_198380 (928aa) | maker-scaffold10x_919_pilon-snap-gene-0.104 | 675 | 17.00 | Argonaute, linker 1 domain (IPR014811)<br>PAZ domain (IPR003100)<br>Ribonuclease H-like domain (IPR012337)<br>Piwi domain (IPR003165) | **Piwi Magnesium Ion Interacting Residues**<br>GADVTHP…YRDGV | |
| **Argonaute-2** | Smp_179320 (932aa) | maker-scaffold10x_145_pilon-augustus-gene-2.246 (BN1106_s1192B000304) | 355 (1024) | 16.8 (28.3) | Protein argonaute, N-terminal (IPR032474)<br>Argonaute, linker 1 domain (IPR014811)<br>PAZ domain (IPR003100)<br>Argonaute linker 2 domain (IPR032472<br>Protein argonaute, Mid domain (IPR032473)<br>Ribonuclease H-like domain (IPR012337)-<br>Piwi domain (IPR003165) | **Piwi Magnesium Ion Interacting Residues**<br>GADVTHP…YRDGV…YYAHL | |
| **Argonaute-3** | Smp_102690 (854aa) | maker-scaffold10x_979_pilon-snap-gene-0.4 | 898 | 36.60 | Argonaute, linker 1 domain (IPR014811), Ribonuclease H-like domain (IPR012337)<br>Piwi domain (IPR003165) | **Piwi Magnesium Ion Interacting Residues**<br>GADVTHP…YRDGV…YYSHL | |
| **Tudor-SN** | Smp_246840 (992aa) | maker-scaffold10x_485_pilon-snap-gene-0.80 | 1098 | 50.40 | Staphylococcal nuclease (SNase-like), OB-fold (IPR016071), | | |
| | Smp_246850 (378aa) | maker-scaffold10x_1598_pilon-snap-gene-0.1 (BN1106_s375B000233) | 1220 (518) | 8.0 (22.22) | Staphylococcal nuclease (SNase-like), OB-fold (IPR016071), Tudor domain (IPR002999) | | |
| **FMR1** | Smp_099630 (598aa) | BN1106_s2932B000172 | 690 | 20.40 | K Homology domain (IPR004087)<br>K Homology domain, type 1 (IPR004088) | | |
| | | maker-scaffold10x_2403_pilon-snap-gene-0.22 (BN1106_s6067B000089) | 1015 (460) | 24.1 (43.7) | K Homology domain, type 1 (IPR004088) | | |
| | Smp_058170 (827aa) | maker-scaffold10x_142_pilon-snap-gene-0.83 | 812 | 44.70 | K Homology domain, type 1 (IPR004088) | | |

| | | | | | | |
|---|---|---|---|---|---|---|
| | | snap_masked-scaffold10x_1280_pilon-processed-gene-0.18/snap_masked-scaffold10x_1280_pilon-processed-gene-0.19 | 569 | 15.50 | K Homology domain, type 1 (IPR004088) | |
| SID-1 | Smp_152020 (933aa) | maker-scaffold10x_566_pilon-snap-gene-0.12 | 864 | 44.80 | PF13965 (SID-1_RNA_chan) | |
| Exportin-1 | Smp_124820 (1075aa) | maker-scaffold10x_1096_pilon-snap-gene-0.68 | 1211 | 72.30 | Armadillo-type fold (IPR016024)<br>Armadillo-like helical (IPR011989)<br>Importin-beta, N-terminal domain (IPR001494)<br>Exportin-1/Importin-beta-like (IPR013598)<br>CRM1 C-terminal domain (IPR014877) | |
| Exportin-2 | Smp_128050 (1049aa) | maker-scaffold10x_1061_pilon-snap-gene-0.108 | 1061 | 60.00 | Armadillo-type fold (IPR016024)<br>Armadillo-like helical (IPR011989)<br>Importin-beta, N-terminal domain (IPR001494)<br>Exportin/Importin, Cse1-like (IPR013713)<br>Exportin/Importin, Cse1-like (IPR013713)<br>CAS/CSE, C-terminal (IPR005043) | |
| Exportin-5 | Smp_152800 (1286aa) | snap_masked-scaffold10x_1656_pilon-processed-gene-0.32 | 786 | 25.10 | Armadillo-type fold (IPR016024)<br>Armadillo-like helical (IPR011989) | |
| ERI1 | Smp_029850 (562aa) | maker-scaffold10x_1695_pilon-pred_gff_StringTie-gene-0.47 | 788 | 38.70 | Ribonuclease H-like domain (IPR012337)<br>Exonuclease, RNase T/DNA polymerase III (IPR013520) | Nuclease D…E…D…H…D |
| ERI2 | Smp_051420 (232aa) | maker-scaffold10x_48_pilon-snap-gene-0.20 | 240 | 57.45 | Ribonuclease H-like domain (IPR012337)<br>Exonuclease, RNase T/DNA polymerase III (IPR013520) | Nuclease D…H…D |
| ERI3* | Smp_157760 (191aa) | BN1106_s1255B000315.1 | 211 | 54.30 | Ribonuclease H-like domain (IPR012337)<br>Exonuclease, RNase T/DNA polymerase III (IPR013520) | Nuclease D…E…D…H…D |
| | | BN1106_s1255B000315.2 | 85 | 55.95 | Ribonuclease H-like domain (IPR012337)<br>Exonuclease, RNase T/DNA polymerase III (IPR013520) | Nuclease H…D |
| SMG2 | Smp_061590 (1325aa) | maker-scaffold10x_513_pilon-snap-gene-1.141 | 1336 | 74.10 | RNA helicase UPF1, UPF2-interacting domain (IPR018999)<br>P-loop containing nucleoside triphosphate hydrolase (IPR027417) | |
| SMG4 | Smp_102240 (560aa) | maker-scaffold10x_637_pilon-snap-gene-0.146 | 456 | 12.90 | Regulator of nonsense-mediated decay, UPF3 (IPR005120)<br>Nucleotide-binding alpha-beta plait domain (IPR012677) | |
| SMG5 | Smp_169530 (1479aa) | maker-scaffold10x_503_pilon-snap-gene-0.67 | 1658 | 31.50 | Tetratricopeptide-like helical domain (IPR011990)<br>DNA/RNA-binding domain, Est1-type (IPR018834) | |
| SMG6 | Smp_131650 (1409aa) | maker-scaffold10x_102_pilon-augustus-gene-0.93 (BN1106_s2800B000128) | 224 (1381) | 3.3 (54.2) | Tetratricopeptide-like helical domain (IPR011990)<br>Telomerase activating protein Est1 (IPR019458)<br>DNA/RNA-binding domain, Est1-type (IPR018834)<br>PIN domain-like (IPR029060)<br>PIN domain (IPR002716) | |

*These predicted genes are absent from new genome assembly, however PCR data have confirmed their presence and expression in juvenile and adult worms

### Supplementary File 1

#### *Drosha Sequences*
Q9NRR4.2 (NCBI)
XP_010815134.1 (NCBI)
NP_477436.1 (NCBI)
CE29039 (Wormbase)
BM25607 (Wormbase)
CN03400 (Wormbase)
CBP41277 (Wormbase)
JA61114 (Wormbase)
RP42472 (Wormbase)
OVP08524 (Wormbase)
Sjp_0048900 (Wormbase Parasite)
Smp_142510 (Wormbase Parasite)
Ancylostoma caninum (Dalzell et al. 2011)
Ascaris suum (Dalzell et al. 2011)
Meloidogyne hapla (Dalzell et al. 2011)
Meloidogyne incognita (Dalzell et al. 2011)
Pristionchus pacificus (Dalzell et al. 2011)
Trichinella spiralis (Dalzell et al. 2011)

#### *DGCR8 Sequences*
NP_073557.3 (NCBI)
NP_001094674.1 (NCBI)
NP_651879.1 (NCBI)
Smp_087220 (Wormbase Parasite)
Sjp_0013270 (Wormbase Parasite)
CE48538 PASH-1 (Wormbase)
BM38243 PASH-1 (Wormbase)
CBP43006 PASH-1 (Wormbase)
JA66543 PASH-1 (Wormbase)
RP39638 PASH-1 (Wormbase)
OVP13372 PASH-1 (Wormbase)
PP03672 PASH-1 (Wormbase)
Caenorhabditis brenneri (Dalzell et al. 2011)
Haemonchus contortus (Dalzell et al. 2011)
Meloidogyne hapla (Dalzell et al. 2011)
Meloidogyne incognita (Dalzell et al. 2011)

#### *Dicer Sequences*
NP_001278557.1 (NCBI)
AAR26432.1 (NCBI)
NP_524453.1 (NCBI)
NP_523778.2 (NCBI)
EKC26346.1 (NCBI)
Smp_169750 (Wormbase Parasite)
Smp_033600 (Wormbase Parasite)
Sjp_0069770 (Wormbase Parasite)
Sjp_0043700 (Wormbase Parasite)
CE47418 DCR-1 (Wormbase)
BM36861 DCR-1 (Wormbase)
CN04613 DCR-1 (Wormbase)
CBP41384 DCR-1 (Wormbase)
JA44861 DCR-1 (Wormbase)
OVP14022 DCR-1 (Wormbase)
Ancylostoma caninum (Dalzell et al. 2011)
Ascaris suum (Dalzell et al. 2011)
Meloidogyne hapla (Dalzell et al. 2011)
Meloidogyne incognita (Dalzell et al. 2011)
Oesophagostomum dentatum (Dalzell et al. 2011)
Pristionchus pacificus (Dalzell et al. 2011)
Trichinella spiralis (Dalzell et al. 2011)

#### *DRH-1 Sequences*
NP_055129.2 (NCBI)
XP_002689526.3 (NCBI)
EKC38303.1 (NCBI)
CE16988 (Wormbase)
BM40905 (Wormbase)
CN02053 (Wormbase)
CBP38705 (Wormbase)
JA40499 (Wormbase)
RP41755 (Wormbase)
Ancylostoma caninum (Dalzell et al. 2011)
Ascaris suum (Dalzell et al. 2011)
Haemonchus contortus (Dalzell et al. 2011)
Meloidogyne hapla (Dalzell et al. 2011)
Meloidogyne incognita (Dalzell et al. 2011)
Trichinella spiralis (Dalzell et al. 2011)

#### *DRH-3 Sequences*
CE36120 (Wormbase)
BM45565 (Wormbase)
CBP46809 (Wormbase)
JA52630 (Wormbase)
RP48790 (Wormbase)
OVP14314 (Wormbase)
PP30574 (Wormbase)
Ancylostoma caninum (Dalzell et al. 2011)
Ascaris suum (Dalzell et al. 2011)
Caenorhabditis brenneri (Dalzell et al. 2011)
Haemonchus contortus (Dalzell et al. 2011)
Meloidogyne hapla (Dalzell et al. 2011)
Meloidogyne incognita (Dalzell et al. 2011)
Oesophagostomum dentatum (Dalzell et al. 2011)
Trichinella spiralis (Dalzell et al. 2011)

#### *RDE-4 Sequences*
CE00630 (Wormbase)
BM23272 (Wormbase)
CBP08423 (Wormbase)
JA48137 (Wormbase)
RP14524 (Wormbase)
PP50570 (Wormbase)
Ancylostoma caninum (Dalzell et al. 2011)
Caenorhabditis brenneri (Dalzell et al. 2011)

#### *RRF-1 Sequences*
CE27141 (Wormbase)
CBP37456 (Wormbase)
RP46078 (Wormbase)
Ascaris suum (Dalzell et al. 2011)
Caenorhabditis brenneri (Dalzell et al. 2011)

#### *Argonaute-like Sequences*
NP_036331.1 (NCBI)
NP_036286.2 (NCBI)
NP_079128.2 (NCBI)
NP_060099.2 (NCBI)
NP_001192828.1 (NCBI)
AAS21301.1 (NCBI)
AAR12162.2 (NCBI)
NP_725341.1 (NCBI)
NP_648775.1 (NCBI)
EKC35067.1 (NCBI)
Smp_102690 (Wormbase Parasite)
Smp_198380 (Wormbase Parasite)
Smp_179320 (Wormbase Parasite)
Sjp_0044720 (Wormbase Parasite)
Sjp_0045200 (Wormbase Parasite)
Sjp_0103990 (Wormbase Parasite)
CE31525 (Wormbase)
CE32063 (Wormbase)
CE18277 (Wormbase)
CE01117 (Wormbase)
CN03079 (Wormbase)
CBP07776 (Wormbase)
CBP40745 (Wormbase)
JA63108 (Wormbase)
RP37764 (Wormbase)
PP22070 (Wormbase)
Ancylostoma caninum (Dalzell et. al 2011)
Ascaris suum (Dalzell et. al 2011)
Brugia malayi (Dalzell et. al 2011)
Haemonchus contortus (Dalzell et. al 2011)
Meloidogyne hapla (Dalzell et. al 2011)
Meloidogyne incognita (Dalzell et. al 2011)
Oesophagostomum dentatum (Dalzell et. al 2011)
Pristionchus pacificus (Dalzell et. al 2011)
Trichinella spiralis (Dalzell et. al 2011)
Ascaris suum (Dalzell et. al 2011)
Haemonchus contortus (Dalzell et. al 2011)
Meloidogyne hapla (Dalzell et. al 2011)
Meloidogyne incognita (Dalzell et. al 2011)
Trichinella spiralis (Dalzell et. al 2011)

#### *PIWI-Clade AGO Sequences*
AAC97371.2 (NCBI)
BAC81342.1 (NCBI)
BAC81343.1 (NCBI)
NP_689644.2 (NCBI)
NP_001302505.1 (NCBI)
XP_015320079.1 (NCBI)
NP_001306825.1 (NCBI)
XP_015321929.1 (NCBI)
AGL81535.1 (NCBI)
CAA64320.1 (NCBI)
EKC35279.1 (NCBI)
EKC29295.1 (NCBI)
CE30106 ERGO-1 (Wormbase)
CBP27166 ERGO-1 (Wormbase)
JA40180 ERGO-1 (Wormbase)
RP28315 ERGO-1 (Wormbase)
OVP00965 ERGO-1 (Wormbase)
Caenorhabditis brenneri (Dalzell et al. 2011)
Trichinella spiralis (Dalzell et al. 2011)
CE09083 (Wormbase)
CE06748 (Wormbase)
CBP09000 (Wormbase)
Ancylostoma caninum (Dalzell et al. 2011)
Caenorhabditis brenneri (Dalzell et al. 2011)
Caenorhabditis japonica (Dalzell et al. 2011)
Pristionchus pacificus (Dalzell et al. 2011)
Haemonchus contortus (Dalzell et al. 2011)
Oesophagostomum dentatum (Dalzell et al. 2011)

#### *Helminth-only AGO Sequences*
CE06244 (Wormbase)
CE17921 (Wormbase)
CE48028 (Wormbase)
CE20101 (Wormbase)
CE22246 (Wormbase)
CN20858 (Wormbase)
CN25357 (Wormbase)
CN21626 (Wormbase)
JA14615 (Wormbase)
RP04583 (Wormbase)
PP44473 (Wormbase)
Ancylostoma caninum (Dalzell et. al 2011)
Ascaris suum (Dalzell et. al 2011)
Brugia malayi (Dalzell et. al 2011)
Haemonchus contortus (Dalzell et. al 2011)
Meloidogyne hapla (Dalzell et. al 2011)
Meloidogyne incognita (Dalzell et. al 2011)
Oesophagostomum dentatum (Dalzell et. al 2011)
Pristionchus pacificus (Dalzell et. al 2011)
Ancylostoma caninum (Dalzell et. al 2011)
Ascaris suum (Dalzell et. al 2011)
Brugia malayi (Dalzell et. al 2011)
Haemonchus contortus (Dalzell et. al 2011)
Meloidogyne hapla (Dalzell et. al 2011)
Meloidogyne incognita (Dalzell et. al 2011)
Oesophagostomum dentatum (Dalzell et. al 2011)
Ancylostoma caninum (Dalzell et. al 2011)
Ascaris suum (Dalzell et. al 2011)
Haemonchus contortus (Dalzell et. al 2011)
Oesophagostomum dentatum (Dalzell et. al 2011)
Ancylostoma caninum (Dalzell et. al 2011)
Meloidogyne incognita (Dalzell et. al 2011)
Oesophagostomum dentatum (Dalzell et. al 2011)
Pristionchus pacificus (Dalzell et. al 2011)
CE12164 (Wormbase)
CE25906 (Wormbase)
CBP37437 (Wormbase)
CBP39116 (Wormbase)
JA07867 (Wormbase)
RP46812 (Wormbase)
Ancylostoma caninum (Dalzell et. al 2011)
Ascaris suum (Dalzell et. al 2011)
Caenorhabditis brenneri (Dalzell et. al 2011)
Caenorhabditis brenneri (Dalzell et. al 2011)
Caenorhabditis remanei (Dalzell et. al 2011)
Oesophagostomum dentatum (Dalzell et. al 2011)
CE28243 (Wormbase)
BM38004 (Wormbase)
CN02916 (Wormbase)
CBP45899 (Wormbase)
JA45228 (Wormbase)
RP45359 (Wormbase)
Ancylostoma caninum (Dalzell et. al 2011)
Ascaris suum (Dalzell et. al 2011)
Haemonchus contortus (Dalzell et. al 2011)
Pristionchus pacificus (Dalzell et. al 2011)

#### *TSN-1 Sequences*
AAH17180.3 (NCBI)
AAI04505.1 (NCBI)
NP_612021.1 (NCBI)
Smp_246840 (Wormbase Parasite)
Smp_246850 (Wormbase Parasite)
CE02626 (Wormbase)
BM43652 TSN-1 (Wormbase)
CBP01117 (Wormbase)
JA55899 (Wormbase)
RP14297 (Wormbase)
OVP12598 (Wormbase)
PP14068 (Wormbase)
Caenorhabditis brenneri (Dalzell et al. 2011)
Meloidogyne hapla (Dalzell et al. 2011)
Meloidogyne incognita (Dalzell et al. 2011)
Oesophagostomum dentatum (Dalzell et al. 2011)
Trichinella spiralis (Dalzell et al. 2011)

#### *VIG-1 Sequences*
NP_523572.1 (NCBI)
CE11254 (Wormbase)
BM36903 (Wormbase)
CN19739 (Wormbase)
CBP07521 (Wormbase)
JA11839 (Wormbase)
RP37119 (Wormbase)
OVP08750 (Wormbase)
PP06425 (Wormbase)
Ancylostoma caninum (Dalzell et al. 2011)
Ascaris suum (Dalzell et al. 2011)
Oesophagostomum dentatum (Dalzell et al. 2011)

#### *ADAR Sequences*
P55265.4 (NCBI)
XP_015318055.1 (NCBI)
NP_569940.2 (NCBI)
CE32459 (Wormbase)
CBP39236 (Wormbase)
RP07872 (Wormbase)
Ancylostoma caninum (Dalzell et. al 2011)
Ascaris suum (Dalzell et. al 2011)
Brugia malayi (Dalzell et. al 2011)
Caenorhabditis brenneri (Dalzell et. al 2011)
Ancylostoma caninum (Dalzell et. al 2011)
Ascaris suum (Dalzell et. al 2011)
Brugia malayi (Dalzell et. al 2011)
Caenorhabditis brenneri (Dalzell et. al 2011)
Haemonchus contortus (Dalzell et. al 2011)
Oesophagostomum dentatum (Dalzell et. al 2011)

#### *ADAR-2 Sequences*
P78563.1 (NCBI)
XP_005202097.1 (NCBI)
AAF63702.1 (NCBI)
CE25120 (Wormbase)
CN05541 (Wormbase)
CBP24602 (Wormbase)
JA47475 (Wormbase)
RP33924 (Wormbase)
PP29926 (Wormbase)
Ancylostoma caninum (Dalzell et. al 2011)
Ascaris suum (Dalzell et. al 2011)
Haemonchus contortus (Dalzell et. al 2011)
Meloidogyne hapla (Dalzell et. al 2011)
Oesophagostomum dentatum (Dalzell et. al 2011)

#### *ERI1 Sequences*
NP_699163.2 (NCBI)
NP_001033281.1 (NCBI)
XP_011426949.1 (NCBI)
CE17214 (Wormbase)
Sjp_0057290 (Wormbase Parasite)
Smp_051420.1 (Wormbase Parasite)
Smp_157760 (Wormbase Parasite)
Ascaris suum (Dalzell et. al 2011)
BM22822 (Wormbase)
Caenorhabditis brenneri (Dalzell et. al 2011)
CBP17865 (Wormbase)
JA63029 (Wormbase)
RP30882 (Wormbase)
Haemonchus contortus (Dalzell et. al 2011)
Meloidogyne hapla (Dalzell et. al 2011)
Meloidogyne incognita (Dalzell et. al 2011)
Oesophagostomum dentatum (Dalzell et. al 2011)
PP:PP45833 Pristionchus pacificus (Dalzell et. al 2011)
Trichinella spiralis (Dalzell et. al 2011)

#### *ERI-3 Sequences*
CE32072 (Wormbase)
CBP45144 (Wormbase)
RP30649 (Wormbase)

#### *ERI5 Sequences*
CE39434 (Wormbase)
CN20393 (Wormbase)
CBP45144 (Wormbase)
JA61238 (Wormbase)
RP23017 (Wormbase)

#### *ERI6 Sequences*
CE08658 (Wormbase)
Caenorhabditis brenneri (Dalzell et. al 2011)
Caenorhabditis briggsae (Dalzell et. al 2011)
Caenorhabditis japonica (Dalzell et. al 2011)
Caenorhabditis remanei (Dalzell et. al 2011)

#### *ERI7 Sequences*
CE36108 (Wormbase)
Caenorhabditis brenneri (Dalzell et. al 2011)
Caenorhabditis briggsae (Dalzell et. al 2011)
Caenorhabditis japonica (Dalzell et. al 2011)
Caenorhabditis remanei (Dalzell et. al 2011)

#### *lin-15b Sequences*
CE15381 (Wormbase)
CBP44496 (Wormbase)
JA63349 (Wormbase)
Caenorhabditis brenneri (Dalzell et. al 2011)
Caenorhabditis remanei (Dalzell et. al 2011)

#### *XRN1 Sequences*
AAH39314.1 (NCBI)
XP_005201950.1 (NCBI)
NP_523408.2 (NCBI)
XP_011444428.1 (NCBI)
Smp_140420 (Wormbase Parasite)
Sjp_0051080 (Wormbase Parasite)
CE42924 (Wormbase)
BM45312 (Wormbase)
CN18416 (Wormbase)
CBP41142 (Wormbase)
JA65206 (Wormbase)
RP29844 (Wormbase)
OVP05624 (Wormbase)
PP43664 (Wormbase)
Ascaris suum (Dalzell et. al 2011)
Haemonchus contortus (Dalzell et. al 2011)
Oesophagostomum dentatum (Dalzell et. al 2011)
Trichinella spiralis (Dalzell et. al 2011)

#### *XRN2 Sequences*
NP_001304889.1 (NCBI)
XP_010809536.1 (NCBI)
NP_609082.1 (NCBI)
XP_011451785.1 (NCBI)
Smp_133300 (Wormbase Parasite)
Sjp_0075730 (Wormbase Parasite)
CE42702 (Wormbase)
CBP30614 (Wormbase)
JA59509 (Wormbase)
RP35486 (Wormbase)
OVP13756 (Wormbase)
PP42135 (Wormbase)
Ancylostoma caninum (Dalzell et. al 2011)
Ascaris suum (Dalzell et. al 2011)
Brugia malayi (Dalzell et. al 2011)
Caenorhabditis brenneri (Dalzell et. al 2011)
Haemonchus contortus (Dalzell et. al 2011)
Meloidigyne hapla (Dalzell et. al 2011)
Meloidogyne incognita (Dalzell et. al 2011)
Oesophagostomum dentatum (Dalzell et. al 2011)
Trichinella spiralis (Dalzell et. al 2011)

#### *RSD-2 Sequences*
CE40653 (Wormbase)
CBP41205 (Wormbase)
JA45044 (Wormbase)
Caenorhabditis remanei (Dalzell et. al 2011)

#### *RSD-3 Sequences*
Q14677.1 (NCBI)
NP_001098887.1 (NCBI)
EKC32147.1 (NCBI)
Sjp_0064420 (Wormbase Parasite)
CE03062 (Wormbase)
BM19311 (Wormbase)
CBP34856 (Wormbase)
JA44423 (Wormbase)
RP30343 (Wormbase)
OVP04093 (Wormbase)
PP29990 (Wormbase)
Ancylostoma caninum (Dalzell et. al 2011)
Ascaris suum (Dalzell et. al 2011)
Caenorhabditis brenneri (Dalzell et. al 2011)
Haemonchus contortus (Dalzell et. al 2011)
Meloidogyne hapla (Dalzell et. al 2011)
Meloidogyne incognita (Dalzell et. al 2011)
Oesophagostomum dentatum (Dalzell et. al 2011)
Trichinella spiralis (Dalzell et. al 2011)

#### *RSD-6 Sequences*
B5MCY1.1 (NCBI)
XP_015329037.1 (NCBI)
CE32384 (Wormbase)
CN04313 (Wormbase)
CBP14864 (Wormbase)
JA42867 (Wormbase)
RP29689 (Wormbase)
Pristionchus pacificus (Dalzell et. al 2011)

#### *SID-1 Sequences*
AAI17223.1 (NCBI)
XP_005201330.1 (NCBI)
Smp_152020 (Wormbase Parasite)
Sjp_0101810/Sjp_0067130 (Wormbase Parasite)
CE30331 SID1 (Wormbase)
BM45574 SID1 (Wormbase)
CN02065 (Wormbase)
CBP22114 (Wormbase)
JA60059 (Wormbase)
RP37195 (Wormbase)
OVP05450 (Wormbase)
Haemonchus contortus (Dalzell et. al 2011)
Oesophogostamum dentatum (Dalzell et. al 2011)

#### *SID-2 Sequences*
CE33908 (Wormbase)
CN00799 (Wormbase)
CBP19178 (Wormbase)
JA02006 (Wormbase)
RP38789 (Wormbase)

#### *EGO-1 Sequences*
CE27140 (Wormbase)
CN2005 (Wormbase)
CBP37945 (Wormbase)
JA41806 (Wormbase)
Ascaris suum (Dalzell et. al 2011)
Brugia malayi (Dalzell et. al 2011)
Meloidogyne hapla (Dalzell et. al 2011)
Meloidogyne incognita (Dalzell et. al 2011)
Oesophagostomum dentatum (Dalzell et. al 2011)

#### *MUT-7 Sequences*
NP_060290.3 (NCBI)
XP_015321064.1 (NCBI)
XP_011434475.1 (NCBI)
NP_610094.1 (NCBI)
CE00370 (Wormbase)
CN38432 (Wormbase)
CBP46637 (Wormbase)
JA59535 (Wormbase)
RP29673 (Wormbase)
Ancylostoma caninum (Dalzell et. al 2011)
Ascaris suum (Dalzell et. al 2011)
Brugia malayi (Dalzell et. al 2011)
Haemonchus contortus (Dalzell et. al 2011)
Meloidogyne hapla (Dalzell et. al 2011)
Oesophagostomum dentatum (Dalzell et. al 2011)
Pristionchus pacificus (Dalzell et. al 2011)
Trichinella spiralis (Dalzell et. al 2011)

#### *RDE-2 Sequences*
CE37005 (Wormbase)
CBP43782 (Wormbase)
JA47859 (Wormbase)
RP28943 (Wormbase)
Caenorhabditis brenneri (Dalzell et. al 2011)

#### *SMG-2 Sequences*
Q92900.2 (NCBI)
DAA28309.1 (NCBI)
EKC21342.1 (NCBI)
NP_572767.1 (NCBI)
Sjp_0010660 (Wormbase Parasite)
Smp_061590.1 (Wormbase Parasite)
CE28367 (Wormbase)
BM43052 (Wormbase)
CN24885 (Wormbase)
CBP02073 (Wormbase)
JA66560 (Wormbase)
RP46643 (Wormbase)
OVP14320 (Wormbase)
PP44786 (Wormbase)
Ancylostoma caninum (Dalzell et. al 2011)
Ascaris suum (Dalzell et. al 2011)
Haemonchus contortus (Dalzell et. al 2011)
Meloidogyne hapla (Dalzell et. al 2011)
Meloidogyne incognita (Dalzell et. al 2011)
Oesophagostomum dentatum (Dalzell et. al 2011)
Trichinella spiralis (Dalzell et. al 2011)

#### *SMG-5 Sequences*
BAC53623.1 (NCBI)
NP_001077148.1 (NCBI)
Sjp_0091360 (Wormbase Parasite)
Smp_169530 (Wormbase Parasite)
CE25134 (Wormbase)
CN05132 (Wormbase)
CBP14967 (Wormbase)
JA21381 (Wormbase)
RP21962 (Wormbase)

#### *SMG-6 Sequences*
Q86US8.2 (NCBI)
NP_001192756.1 (NCBI)
EKC33850.1 (NCBI)
Smp_131650 (Wormbase Parasite)
CE32984 (Wormbase)
BM34749 (Wormbase)
CBP43361 (Wormbase)
JA01269 (Wormbase)
OVP04827 (Wormbase)
Ancylostoma caninum (Dalzell et. al 2011)
Ascaris suum (Dalzell et. al 2011)
Caenorhabditis brenneri (Dalzell et. al 2011)
VFLRSCILVTGDRGLTIKAIGNNFPCRGIINFAKWMGV
Haemonchus contortus (Dalzell et. al 2011)
Meloidogyne hapla (Dalzell et. al 2011)
Meloidogyne incognita (Dalzell et. al 2011)
Oesophagostomum dentatum (Dalzell et. al 2011)

Pristionchus pacificus (Dalzell et. al 2011)
Trichinella spiralis (Dalzell et. al 2011)

#### *CID-1 Sequences*

NP_001009881.1 (NCBI)
XP_010801816.1 (NCBI)
XP_011517316.1 (NCBI)
XP_010806440.1 (NCBI)
CE02015 (Wormbase)
CN15335 (Wormbase)
CBP20090 (Wormbase)
JA53700 (Wormbase)
RP42885 (Wormbase)
OVP03099 (Wormbase)

#### *EKL-1 Sequences*

CE05687 (Wormbase)
BM41207 (Wormbase)
CBP37258 (Wormbase)
JA60996 (Wormbase)
RP28013 (Wormbase)
PP41586 (Wormbase)
Ancylostoma caninum (Dalzell et. al 2011)
Caenorhabditis brenneri (Dalzell et. al 2011)
Haemonchus contortus (Dalzell et. al 2011)
Meloidogyne hapla (Dalzell et. al 2011)
Meloidogyne incognita (Dalzell et. al 2011)
Oesophagostomum dentatum (Dalzell et. al 2011)

#### *EKL-4 Sequences*

NP_001029195.1 (NCBI)
EKC31208.1 (NCBI)
Smp_002160 (Wormbase Parasite)
Sjp_0000610 (Wormbase Parasite)
CE29836 (Wormbase)
BM20893 (Wormbase)
CBP08051 (Wormbase)
JA39535 (Wormbase)
RP44403 (Wormbase)
OVP05333 (Wormbase)
PP48715 (Wormbase)
Ascaris suum (Dalzell et. al 2011)
Caenorhabditis brenneri (Dalzell et. al 2011)
Haemonchus contortus (Dalzell et. al 2011)
Meloidogyne hapla (Dalzell et. al 2011)
Meloidogyne incognita (Dalzell et. al 2011)

#### *EKL-5 Sequences*

CE44333 (Wormbase)
CN17207 (Wormbase)
JA53094 (Wormbase)
RP10702 (Wormbase)
Caenorhabditis briggsae (Dalzell et. al 2011)

#### *EKL-6 Sequences*

NP_078838.1 (NCBI)
XP_019834464.1 (NCBI)
NP_609922.2 (NCBI)
CE51593 (Wormbase)
BM41419 (Wormbase)
CN05593 (Wormbase)
JA59875 (Wormbase)
RP31352 (Wormbase)
OVP10637 (Wormbase)
PP34905 (Wormbase)
Ancylostoma caninum (Dalzell et. al 2011)
Caenorhabditis briggsae (Dalzell et. al 2011)
Haemonchus contortus (Dalzell et. al 2011)

#### *GFL-1 Sequences*

NP_006521.1 (NCBI)
NP_001069833.1 (NCBI)
XP_011427964.1 (NCBI)
AAF52462.2 (NCBI)
Smp_070760 (Wormbase Parasite)
Sjp_0068080 (Wormbase Parasite)
CE12388 (Wormbase)
BM41536 (Wormbase)
CBP15531 (Wormbase)
JA12698 (Wormbase)
RP19851 (Wormbase)
OVP11512 (Wormbase)
Ancylostoma caninum (Dalzell et. al 2011)
Ascaris suum (Dalzell et. al 2011)
Haemonchus contortus (Dalzell et. al 2011)
Meloidogyne incognita (Dalzell et. al 2011)
Oesophagostomum dentatum (Dalzell et. al 2011)
Trichinella spiralis (Dalzell et. al 2011)

#### *MES-2 Sequences*

CAA64955.1 (NCBI)
NP_001095621.1 (NCBI)
NP_524021.2 (NCBI)
Smp_164650 (Wormbase Parasite)
Sjp_0072180 (Wormbase Parasite)
CE28067 (Wormbase)
BM29030 (Wormbase)
CN34880 (Wormbase)
CBP33838 (Wormbase)
JA64995 (Wormbase)
OVP09541 (Wormbase)
Ancylostoma caninum (Dalzell et. al 2011)
Ascaris suum (Dalzell et. al 2011)
Caenorhabditis remanei (Dalzell et. al 2011)
Haemonchus contortus (Dalzell et. al 2011)
Meloidogyne incognita (Dalzell et. al 2011)
Oesophagostomum dentatum (Dalzell et. al 2011)
Trichinella spiralis (Dalzell et. al 2011)

#### *MES-3 Sequences*

CE11046 (Wormbase)
CBP38928 (Wormbase)
JA67604 (Wormbase)
RP41043 (Wormbase)
Caenorhabditis brenneri (Dalzell et. al 2011)

#### *MES-4 Sequences*

CE27781 (Wormbase)
BM24937 (Wormbase)
CBP07413 (Wormbase)
JA55637 (Wormbase)
RP48484 (Wormbase)
OVP14309 (Wormbase)
PP39245 MES-4 (Wormbase)

#### *MES-6 Sequences*

O75530.2 (NCBI)
Q3SZ25.1 (NCBI)
Q26458.1 (NCBI)
Smp_165220 (Wormbase Parasite)
CE24796 (Wormbase)
BM42303 (Wormbase)
CN03316 (Wormbase)
CBP39565 (Wormbase)
JA02512 (Wormbase)
RP31594 (Wormbase)
OVP05048 (Wormbase)
Ancylostoma caninum (Dalzell et. al 2011)
Ascaris suum (Dalzell et. al 2011)
Haemonchus contortus (Dalzell et. al 2011)
Meloidogyne hapla (Dalzell et. al 2011)
Oesophagostomum dentatum (Dalzell et. al 2011)

#### *MUT-2 Sequences*

CE11740 (Wormbase)
CN15799 (Wormbase)
CBP17632 (Wormbase)
JA58183 (Wormbase)
RP24216 (Wormbase)
PP42162 (Wormbase)
Haemonchus contortus (Dalzell et. al 2011)

#### *MUT-14 Sequences*

CE06825 (Wormbase)
CE28975 (Wormbase)
CBP16557 (Wormbase)
RP31627 (Wormbase)
JA05879 (Wormbase)

#### *MUT-15 Sequences*

CE23936 (Wormbase)
CBP46299 (Wormbase)
JA57376 (Wormbase)
RP44547 (Wormbase)

#### *MUT-16 Sequences*

CE40346 (Wormbase)
CBP44329 (Wormbase)
JA63728 (Wormbase)
RP48608 (Wormbase)
Caenorhabditis brenneri (Dalzell et. al 2011)

#### *RHA-1 Sequences*

NP_001348.2 (NCBI)
NP_776461.1 (NCBI)
NP_476641.1 (NCBI)
CE39177 (Wormbase)
BM42381 (Wormbase)
CN00840 (Wormbase)
CBP39763 (Wormbase)
JA40844 (Wormbase)
RP26542 (Wormbase)
Ascaris suum (Dalzell et. al 2011)
Haemonchus contortus (Dalzell et. al 2011)
Meloidogyne hapla (Dalzell et. al 2011)
Meloidogyne incognita (Dalzell et. al 2011)
Trichinella spiralis (Dalzell et. al 2011)

#### *RRF-3 Sequences*

CE45624 (Wormbase)
BM43371 (Wormbase)
CN14535 (Wormbase)
CBP00181 (Wormbase)
JA02852 (Wormbase)
RP37819 (Wormbase)
OVP12943 (Wormbase)
PP41583 (Wormbase)
Ancylostoma caninum (Dalzell et. al 2011)
Ascaris suum (Dalzell et. al 2011)
Haemonchus contortus (Dalzell et. al 2011)
Trichinella spiralis (Dalzell et. al 2011)

#### *ZFP-1 Sequences*

AAT47519.1 (NCBI)
XP_005194335.3 (NCBI)
EKC26968.1 (NCBI)
CE25003 (Wormbase)
BM37915 (Wormbase)
CN00487 (Wormbase)
CBP39359 (Wormbase)
JA66345 (Wormbase)
RP38280 (Wormbase)
PP37542 (Wormbase)
Ascaris suum (Dalzell et. al 2011)
Brugia malayi (Dalzell et. al 2011)
Haemonchus contortus (Dalzell et. al 2011)
Meloidogyne hapla (Dalzell et. al 2011)

**Supplementary File 2**

*fhcam2* and *F. hepatica* cathepsin L (*fhcatL*) dsRNAs were generated as described in main methods (primers listed in Supplementary Table 2). Glass pipettes were pulled using a PC-10 Narishige Puller (3½" glass pipettes; 7 mm drop; Heater 1 setting: 59.1; Heater 2 setting: 58.1) and broken approximately 1 mm from the end to enable liquid to be drawn up. A Drummond "Nanoject" XV Microinjector (Drummond Scientific Company) was used to inject adult liver fluke (those that had regurgitated gut contents) with 18.2 nl of dsRNA (100 ng/µl) or RPMI at seven sites around the worm. Adult worms were maintained in RPMI+ (RPMI with gentamicin (Sigma-Aldrich) diluted 1 in 100) for 18 h before transfer to CS20 (20% chicken serum in RPMI+). Those adults that were soaked in dsRNA were soaked in 5 ml of 100 ng/µl dsRNA for 18 h prior to transfer to CS20. All adults were maintained in 30 ml media for 48 h post dsRNA exposure before being snap frozen for further analysis. All adults were maintained in groups of three.

Adult liver fluke RNA was extracted by lysing single adults in 1 ml of TRIzol® (ThermoFisher Scientific) and following kit instructions. RNA pellets were suspended in 50 µl nuclease free water with 2000 ng of nucleic acid (measured on Nanodrop 2000) DNase treated prior to reverse transcription as described in main methods. qPCRs were run on diluted cDNA (1:10) and data treated in the same way as described in main methods. Protein was extracted using RIPA buffer, as described in McVeigh et al. (2014), although 1 ml extraction buffer was used. Actin (20–33, Sigma Aldrich) was used as control band with analysis carried out as described in main methods.
